# *Listeria monocytogenes* TcyKLMN cystine/cysteine transporter facilitates glutathione synthesis and virulence gene expression

**DOI:** 10.1101/2021.09.07.459368

**Authors:** Moran Brenner, Sivan Friedman, Adi Haber, Ilya Borovok, Nadejda Sigal, Oded Lewinson, Anat A. Herskovits

## Abstract

*Listeria monocytogenes* (*Lm*) is a saprophyte and a human intracellular pathogen. Upon invasion into mammalian cells, it senses multiple metabolic and environmental signals that collectively trigger its transition to the pathogenic state. One of these signals is the tripeptide glutathione, which acts as an allosteric activator of *Lm*’s master virulence regulator, PrfA. While glutathione synthesis by *Lm* was shown to be critical for PrfA activation and virulence gene expression, it remains unclear how this tripeptide is synthesized under changing environments, especially in light of the observation that *Lm* is auxotrophic to one of its precursors, cysteine. Here, we show that the ABC transporter TcyKLMN is a cystine/cysteine importer that supplies cysteine for glutathione synthesis, hence mediating the induction of the virulence genes. Further, we demonstrate that this transporter is negatively regulated by three metabolic regulators: CodY, CymR and CysK, which sense and respond to changing concentrations of branched chain amino acids (BCAA) and cysteine. The data indicate that under low concentrations of BCAA, TcyKLMN is up-regulated, driving the production of glutathione by supplying cysteine, thereby facilitating PrfA activation. These findings provide molecular insight into the coupling of *Lm* metabolism and virulence, connecting BCAA sensing to cysteine uptake and glutathione biosynthesis, as a mechanism that controls virulence gene expression. This study exemplifies how bacterial pathogens sense their intracellular environment and exploit essential metabolites as effectors of virulence.

**Importance:** Bacterial pathogens sense the repertoire of metabolites in the mammalian niche and use this information to shift into a pathogenic state to accomplish successful infection. Glutathione is a virulence-activating signal that is synthesized by *L. monocytogenes* during infection of mammalian cells. In this study, we show that cysteine uptake via TcyKLMN drives glutathione synthesis and virulence gene expression. The data emphasize the intimate cross-regulation between metabolism and virulence in bacterial pathogens.

## Introduction

*Listeria monocytogenes* (*Lm*) is a Gram-positive, facultative, intracellular pathogen and the causative agent of listeriosis, a disease that can lead to severe clinical manifestations in pregnant women, neonates and immunocompromised adults (1). *Lm* is characterized by its intracellular lifestyle, but can also grow outside the host on soil and vegetation, as well as on food products (2). In the mammalian host, *Lm* invades a wide array of cells (phagocytic and non-phagocytic) by expressing specialized proteins, named internalins, that facilitate its internalization (e.g., InlA and InlB) (3). Upon internalization, the bacteria are initially found within a membrane-bound vacuole, from which they escape into the host cell cytosol via the action of several virulence factors - the pore-forming toxin Listeriolysin O (LLO, encoded by the *hly* gene), and the two phospholipases PlcA and PlcB (4–6). In the host cell cytosol, the bacteria utilize host-derived metabolites to support growth (7–10), and using ActA protein hijack the host actin-polymerization machinery to move around the cell and spread from cell to cell (11, 12). All of the above-mentioned virulence factors as well as many others are positively regulated by PrfA, the master virulence regulator of *Lm* (13, 14).

Multiple metabolic and physiological signals are responsible for the transition of *Lm* from the saprophytic to the pathogenic state (15–20). Many of these signals converge at PrfA, and directly or indirectly regulate its transcription, translation and activity. One of these signals is low concentration of branched-chain amino acids (low BCAA), a condition that is found in the intracellular niche. It was previously demonstrated by our lab that *Lm* responds to low BCAA by upregulating the expression of PrfA, which, in turn, activates the virulence genes (15). This response was shown to depend on the global metabolic regulator CodY, which is also a sensor of BCAA, as it directly binds isoleucine (15, 21, 22). Under high-BCAA conditions, CodY was shown to bind isoleucine and in that form to repress the transcription of many metabolic genes, including those involved in BCAA biosynthesis (21–25). While this was considered the main mechanism of CodY regulation, we demonstrated that in *Lm* CodY is also active under low BCAA conditions, i.e., in its isoleucine-unbound form, where it up- and downregulates the transcription of many genes, among them *prfA* (up-regulating its transcription), hence playing a role in the induction of the virulence genes (15, 22). While these findings demonstrated that PrfA expression is essentially linked to the availability of BCAA in the intracellular niche, they further indicated that sensing of the metabolic host cell cytosol environment is key to the regulation of *Lm* virulence.

Another metabolite that was recently shown to affect *Lm* virulence is glutathione. Glutathione is a low-molecular-weight peptide thiol that is highly abundant in the host cell cytosol in its reduced form (GSH), functioning as a redox buffer, antioxidant and enzyme cofactor (26). Glutathione is also synthesized by some bacteria, mainly Gram-negative and few Gram-positive (such as *Lm*), and plays a role in redox homeostasis and bacterial survival under oxidative stress (26). A previous study demonstrated that *Lm*’s glutathione synthetase, GshF, plays a critical role in the induction of the virulence genes during infection (17). GshF is a bifunctional enzyme that catalyzes the two reactions that synthesize the tripeptide glutathione (i.e., L-γ-glutamyl-L-cysteinylglycine) (27). It first ligates the γ-carboxyl group of L-glutamate to L-cysteine, a reaction that is the rate-limiting step of glutathione biosynthesis, and then condenses the product γ-glutamylcysteine with glycine. GSH itself was shown to allosterically bind PrfA, and act as its activating cofactor (28, 29). More specifically, the binding of glutathione to PrfA was shown to cause a conformational change that primes its binding to DNA, as shown for other ligand-binding Crp/Fnr transcription regulators (28, 29).

As indicated, cysteine is one of the building blocks of glutathione, and its rate-limiting precursor, yet, it is not synthesized by *Lm*. *Lm* lacks the ability to reduce sulfate to sulfide, which is required for cysteine biosynthesis, and hence has to import cysteine from the environment (30). Notably, *Lm* is also auxotrophic to methionine (which also has to be imported), and it cannot synthesize cysteine from methionine, as it lacks the transsulfuration pathway that converts methionine to cysteine (9, 30). To date, two transport systems have been shown to be involved in the acquisition of cysteine by *Lm*. The first is the ABC-transporter Lmo0135-0137 (whose substrate binding protein is CtaP), that was shown to support the uptake of free cysteine during growth in a synthetic medium (31). The other system is the OppABCDF transporter, that was shown to import oligopeptides, including cysteine-containing peptides, that were shown to serve as a source of cysteine for glutathione synthesis and PrfA activation (20). While these systems were reported to promote to *Lm* invasion and virulence gene expression in mammalian cells, respectively, their contribution to *Lm* intracellular growth was only partial (20, 31, 32), implying that additional systems are involved in the acquisition of cysteine within the intracellular niche.

In this study, we report on TcyKLMN, another ABC-transporter that is directly involved in cysteine uptake in *Lm*. We show that this transporter imports both cystine and cysteine and plays a role in the activation of *Lm* virulence gene expression by supplying cysteine for glutathione synthesis. Further, we demonstrate that this transporter is regulated by CodY, CymR and CysK, three metabolic factors that act as repressors under nutrient-rich conditions. The findings presented here establish that cysteine import is key to the transition of *Lm* to the pathogenic state, and provide another example of the coupling of metabolism and virulence in bacterial pathogens.

## Results

### Genes associated with cysteine uptake and metabolism modulate virulence gene expression

We previously performed a genetic screen of a mariner transposon mutant library in search of genes that differentially regulate the virulence genes of *Lm* strain 10403S under low BCAA. The screen identified multiple genes that were associated with cysteine uptake and metabolism: *tcyN* (*LMRG_01497*)*, ytmO* (*LMRG_01498*)*, LMRG_01492* and *cysK* (*LMRG_02645*) (Figure 1A and S1A) (33). Interestingly, the first three genes mapped to the *ytmI* operon, which in *B. subtilis* was shown to encode the cystine ABC-transporter, TcyJKLMN (Figure 1A, of note, in *Lm* this operon lacks the *tcyJ* gene) (34). *tcyN* encodes the transporter’s ATP-binding protein, whereas *ytmO* and *LMRG_01492* encode a monooxygenase and an FMN reductase, respectively, whose functions in cystine uptake/metabolism remain elusive. Apart from this locus, the *cysK* gene was also identified, encoding an O-acetylserine thiollyase (which is part of the cysteine synthase complex), which plays a role in cysteine regulation and synthesis (Figure 1A and S1B). In *B. subtilis,* CysK was shown to be a trigger enzyme that acts both as a metabolic enzyme and a transcription regulator, the latter via interaction with CymR, a key transcription regulator in cysteine/methionine metabolism (35). It was shown that CymR and CysK form a complex that negatively regulates the *ytmI* operon by repressing the transcription of its activator YtlI (also called AscR (36, 37), which is encoded upstream to the operon in the opposite direction (a similar gene organization exists in *Lm*, Figure 1A) (38). Although *ytlI* and *cymR* were not identified in our screen, we found homologues in the *Lm* genome (*LMRG_01491* and *LMRG_01455*, respectively), suggesting that the *Lm ytmI* operon is regulated similarly to that of *B. subtilis*.

**Fig 1.**
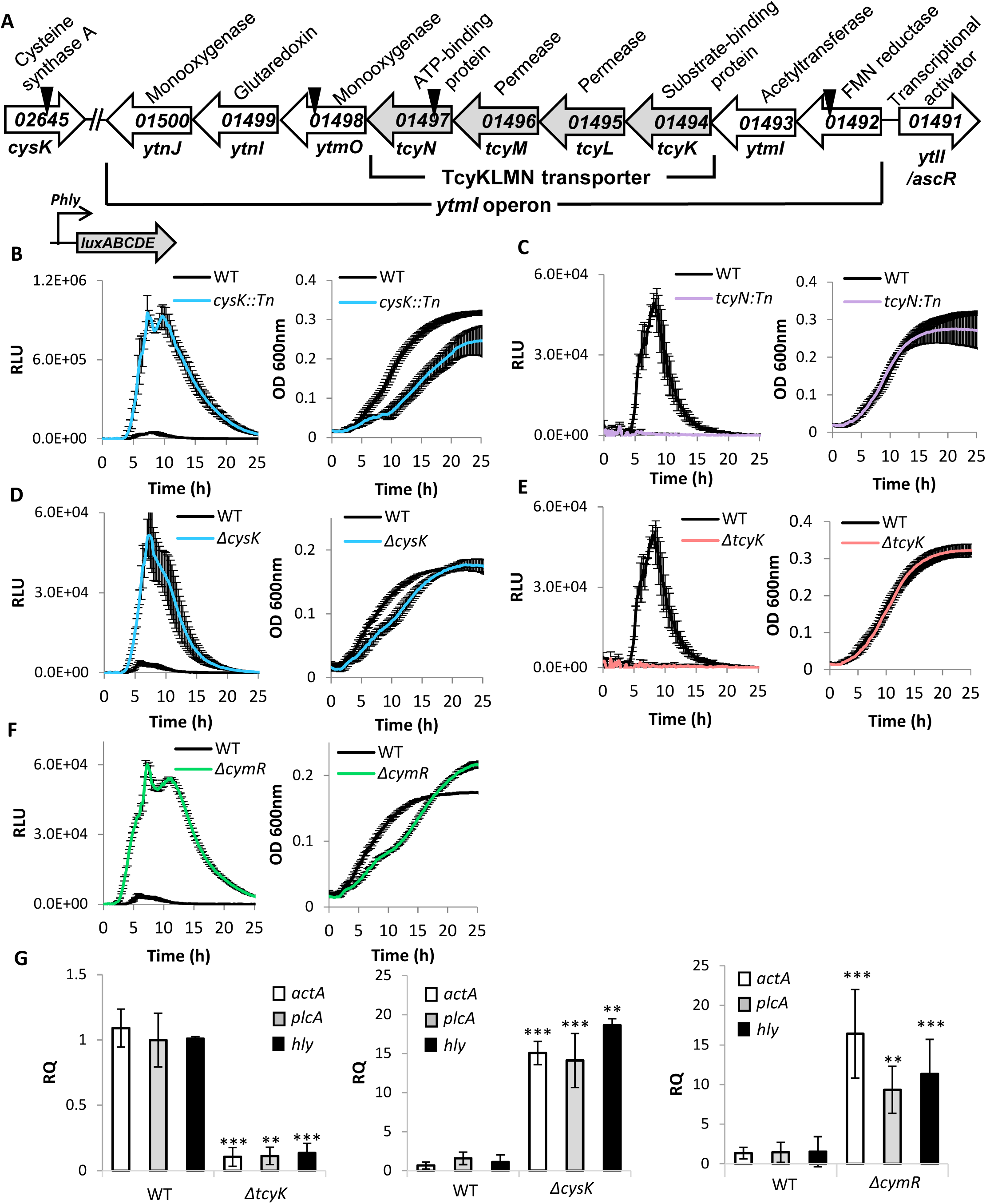
The TcyKLMN transporter and the regulatory factors CysK and CymR play a role in the regulation of *Lm* virulence gene expression. **(A)** A schematic representation of the *ytmI* operon and the genes identified in Friedman *et al*. 2017. Locations of transposon insertions are marked with a triangle. **(B-F)** Luminescence and OD 600nm measurements of bacteria grown in minimal defined media supplemented with low concentrations of branched chain amino acids (LBMM). Relative Luminescence Units (RLU) represent luminescence values normalized to the respective OD 600nm value. OD 600nm values represent growth. The data represent 3 biological replicates. Error bars indicate standard deviation. **(G)** qRT-PCR analysis of *actA*, *plcA* and *hly* transcription levels in indicated bacteria grown in LBMM. mRNA levels were normalized to *rpoD* mRNA and are represented as relative quantity (RQ), relative to mRNA level in WT bacteria. The data represent 3 biological replicates. Error bars indicate standard deviation. Asterisks represent P-values (* = P<0.05, ** = P<0.01, *** = P<0.001, n.s. = non-significant) calculated by Student’s t-test. P-values represent a comparison to the BHI sample.

To investigate the cysteine-associated genes and their role in the regulation of *Lm* virulence, we first examined whether *Lm* strain 10403S is auxotrophic to cysteine and methionine, as some variations have been reported for different *Lm* strains (30). As shown in Figure S2, *Lm* strain 10403S is unable to grow in the absence of cysteine or methionine, and cannot use either of them as a source for the other, hence had to be supplied with both. We next confirmed the phenotypes of the mariner transposon mutants, focusing on *cysK*::*Tn* and *tcyN*::*Tn*, using the same assay that was used in the original screen. Virulence gene expression was monitored using a reporter system that expresses the *luxABCDE* genes under the control of the PrfA-regulated *hly* promoter, which was cloned on the integrative plasmid pPL2 (pPL2-*Phly-lux*). Of note, wild type (WT) bacteria carrying this plasmid grown in a minimal defined medium containing low BCAA (<u>l</u>ow <u>B</u>CAA <u>m</u>inimal <u>m</u>edium, LBMM) display an enhanced luminescence profile, which represents the increased transcription of the *hly* gene [previously shown in (21)]. As mentioned, this upregulation of *hly* transcription in LBMM is completely dependent on CodY, as, under this condition, CodY activates the transcription of PrfA (15, 21). Examining the luminescence profiles of *cysK*::*Tn* and *tcyN*::*Tn* during growth in LBMM, we observed that the mutants differentially affect *Phly-lux* expression in comparison to WT bacteria. While the *cysK::Tn* mutant exhibited an enhanced luminescence profile, *tcyN::Tn* failed to show luminescence signals (Figure 1 B-C) (33). To validate these phenotypes, we generated clean deletion mutants of *cysK* and *tcyK,* the latter encoding the substrate binding protein (SBP) of TcyKLMN (*LMRG_01494*). SBPs of ABC transporters (specifically importers) are key factors that bind the transported substrate extracellularly and facilitate its import into the cell, hence determining the substrate specificity of the transporter (39). Since both *tcyK* and *tcyN* are essential components of TcyKLMN, we chose to delete *tcyK* instead of *tcyN,* to further investigate the transporter’s substrate specificity. As shown in Figure 1 D-E*, ΔcysK* and *ΔtcyK* recapitulated the phenotypes of their corresponding transposon mutants. Of note, the mutants grew similarly to WT bacteria in LBMM and in the rich medium BHI (Figures 1 D-E and Figure S3 A-B). Since in *B. subtilis* CysK was shown to repress *tcyJKLMN* by forming a complex with CymR, we generated a *ΔcymR* mutant of *Lm* and analyzed its luminescence profile during growth in LBMM using the *Phly-lux* reporter system. The data indicated that *ΔcymR* behaves similarly to *ΔcysK,* i.e., exhibits an enhanced luminescence profile, overall demonstrating that CysK and CymR negatively affect the transcription of *hly* (Figure 1F and Figure S3C). To confirm the effects of TcyK, CysK and CymR on the expression of the virulence genes, the transcription levels of *actA*, *plcA* and *hly* (three major virulence genes of *Lm*) were evaluated in *ΔtcyK, ΔcysK* and *ΔcymR* mutants and compared to those in WT bacteria grown in LBMM, using qRT-PCR. As shown in Figure 1G, the transcription level of the virulence genes corroborated the luminescence data, demonstrating a low transcription level in *ΔtcyK* (∼10-fold) and an enhanced transcription level in *ΔcysK* and *ΔcymR* (∼15-fold) in comparison to WT bacteria. These phenotypes were further complemented by introducing a plasmid containing a copy of *cysK* or *tcyK* to the corresponding mutants, demonstrating WT levels of *hly* transcription (Figure S4 and Figure S3 D-E). Taken together, these findings indicated that TcyKLMN, CysK and CymR play a role in the regulation of *Lm* virulence gene expression.

### TcyKLMN is a cystine/cysteine transporter

The transport specificity of ABC transporters that functions as importers is dictated almost exclusively by the binding specificity of their cognate substrate binding proteins (SBP). Therefore, to determine the substrate specificity of *Lm* TcyKLMN, the SBP of the system, TcyK (containing only amino acids 38-286, i.e., without the membrane anchoring domain) was cloned, overexpressed in *E*. *coli* and purified to near homogeneity (Figure S5). We then used isothermal titration calorimetry (ITC) to measure the binding of different substrates to TcyK. We found that TcyK binds cystine (CSSC, the reduced form of cysteine) with a dissociation constant (*K_D_*) of 13 µM and L-cysteine with a *K_D_* of 66.7 µM (Figure 2A). We also tested TcyK for binding of additional amino acids, such as Gln, His, Glu, Thr and Ser, however, none of these were found to be recognized by TcyK (data not shown). To corroborate the ITC findings, we performed growth experiments of *ΔtcyK* and WT bacteria in minimal defined medium containing either cysteine or CSSC as a sole source. Since *Lm* is auxotrophic to cysteine, bacteria with reduced cysteine-import activity should display a growth disadvantage. Indeed, as shown in Figure 2B, when either CSSC or cysteine were added at limited concentrations (<0.5 mM), the growth of *ΔtcyK* was significantly attenuated relative to WT bacteria. These experiments suggest that *Lm* TcyKLMN imports both CSSC and cysteine.

**Fig 2.**
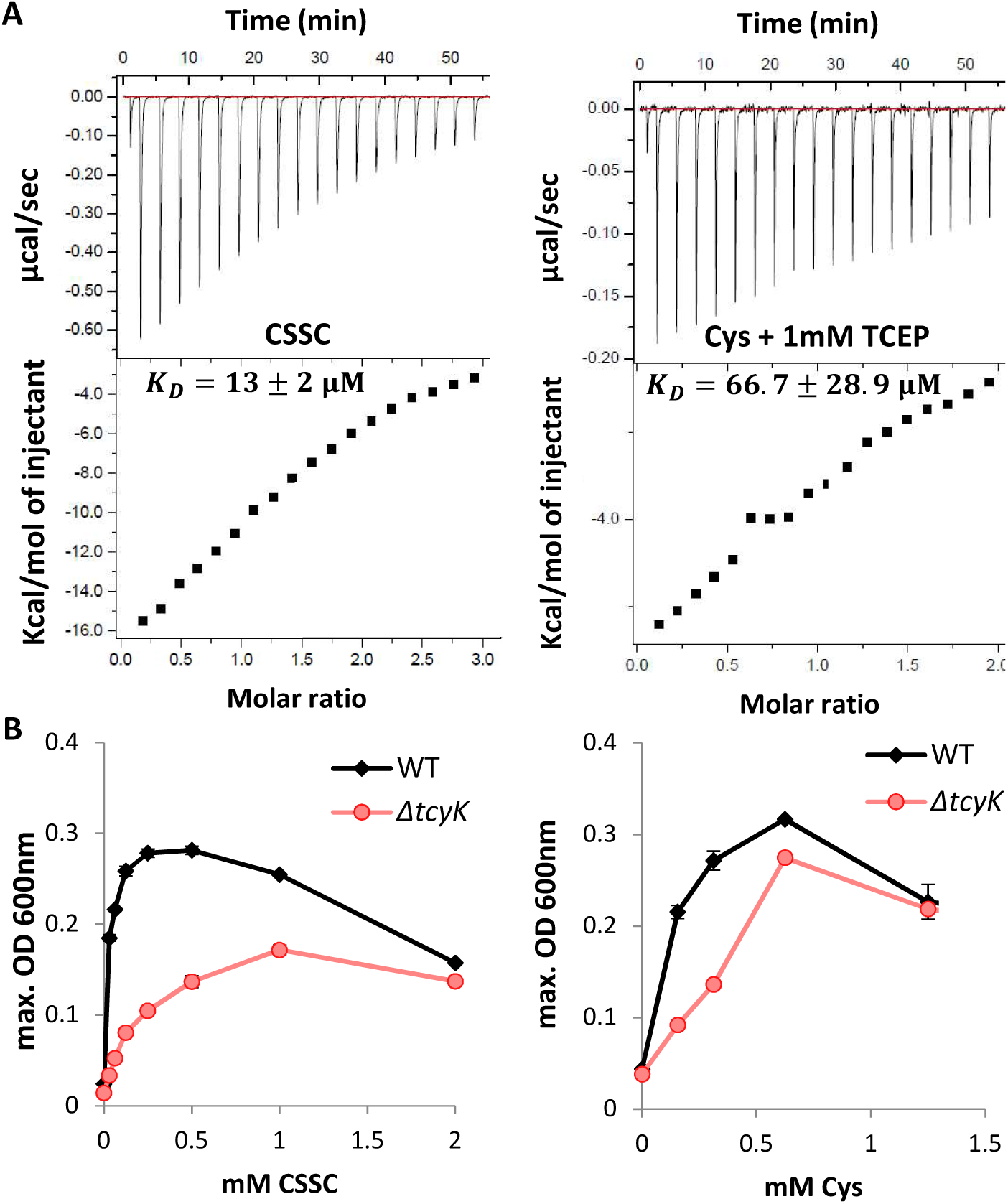
TcyKLMN is a cystine/cysteine transporter. **(A)** Isothermal Titration Calorimetry (ITC) analysis, showing binding of TcyK to cystine (CSSC) or cysteine (Cys). Shown are the consecutive injections of 2 μL aliquots from a 200 μM Cys/CSSC solution into 200 μL of 20 μM TcyK. TCEP was used as a reducing agent when indicated. The upper panels show the calorimetric titration and the lower panels display the integrated injection heat derived from the titrations, for which the best-fit curve was used to calculate the *K_D_*. The experiments were repeated independently 3 times, and the *K_D_* value is presented as a mean ± standard deviation of these 3 experiments. **(B)** Growth of WT *Lm* and *ΔtcyK* bacteria grown in minimal defined media supplemented with different concentrations of cysteine (Cys) or cystine (CSSC) as a sole cysteine source. The data is presented as maximal OD 600nm values, corresponds to bacterial yield. The data represent 2 biological replicates. The experiment was repeated independently twice, with a total of 4 biological replicates. Error bars indicate standard deviation.

### TcyKLMN is negatively regulated by CodY, CysK and CymR

Since TcyKLMN was identified under low BCAA, we next investigated its expression level under this condition. To this end, we compared the transcription levels of *tcyK*, *tcyN* and *ytlI* in WT bacteria grown in LBMM versus BHI. The results demonstrated that the transporter is highly expressed in LBMM (∼100-fold in comparison to BHI, shown by the transcription level of *tcyK* and *tcyN*), and that the YtlI activator gene is also up-regulated (∼10-fold) (Figure 3B). Examining the *ytmI*-*ytlI* intergenic region, we identified two putative CodY binding-sites upstream to the *ytlI* gene (also identified in (22, 40)), suggesting that CodY indirectly regulates TcyKLMN via regulation of the YtlI activator (Figure 3A). To test this hypothesis, we analyzed the transcription levels of *ytlI* and *tcyK* in WT and *ΔcodY* bacteria grown in BHI and LBMM, using qRT-PCR. The experiments confirmed that under nutrient-rich conditions (i.e., in BHI) CodY represses the *ytlI* gene, and consequently *tcyK* (Figure 3C), and further demonstrated that this repression is relieved when BCAA are limited (*i.e.,* in LBMM) (Figure 3D). We next investigated the role of CysK and CymR in the regulation of TcyKLMN (*i.e.,* the *ytmI* operon). Of note, since *Lm* does not assimilate sulfate to sulfide, we assumed that CysK acts as a CymR co-repressor and not as an O-acetylserine thiol-lyase, a function that requires sulfide (Figure S1B). As mentioned, in *B. subtilis* it was shown that CysK and CymR form a complex that represses the TcyJKLMN transporter genes (35). Based on the *B. subtilis* CymR binding-motif (38), we identified two putative CymR binding-sites in the *ytmI*-*ytlI* regulatory region of *Lm*: one upstream the *ytlI* gene and the other upstream the *ytmI* operon (Figure 3A). To examine the potential regulation of this gene locus by CymR and CysK, we first analyzed the transcription levels of *ytlI* and *tcyK* in *ΔcymR* and *ΔcysK* mutants in comparison to WT bacteria grown in BHI and LBMM. The data clearly demonstrated that CymR and CysK act as repressors, down-regulating the transcription of *ytlI* and the *ytmI* operon under both BHI and LBMM conditions (Figure 3 C-D). Since in *B. subtilis* it was shown that CysK and CymR respond to the availability of cysteine, we repeated this experiment using a minimal defined medium containing low concentration of cysteine (LCMM, containing 0.08 mM cysteine). Interestingly, the data demonstrated that under these conditions, the repression of *ytlI* by CymR and CysK was fully relieved, whereas *tcyK* (representative of the transporter genes) was still repressed by these proteins (∼15-fold) (Figure 3E). These findings established that TcyKLMN is negatively regulated, both directly and indirectly, by at least three factors: CodY, CymR and CysK, which collectively respond to changes in BCAA and cysteine levels, and possibly to additional metabolites that affect CymR and CysK. Taken together, the data indicated that under nutrient-rich conditions, all three factors repress the *ytmI* operon, whereas under low BCAA conditions, the operon is up-regulated, albeit, not to its full capacity, as CymR and CysK still repress its transcription to some extent. These findings imply a complex regulation of TcyKLMN in response to different metabolic or environmental cues.

**Fig 3.**
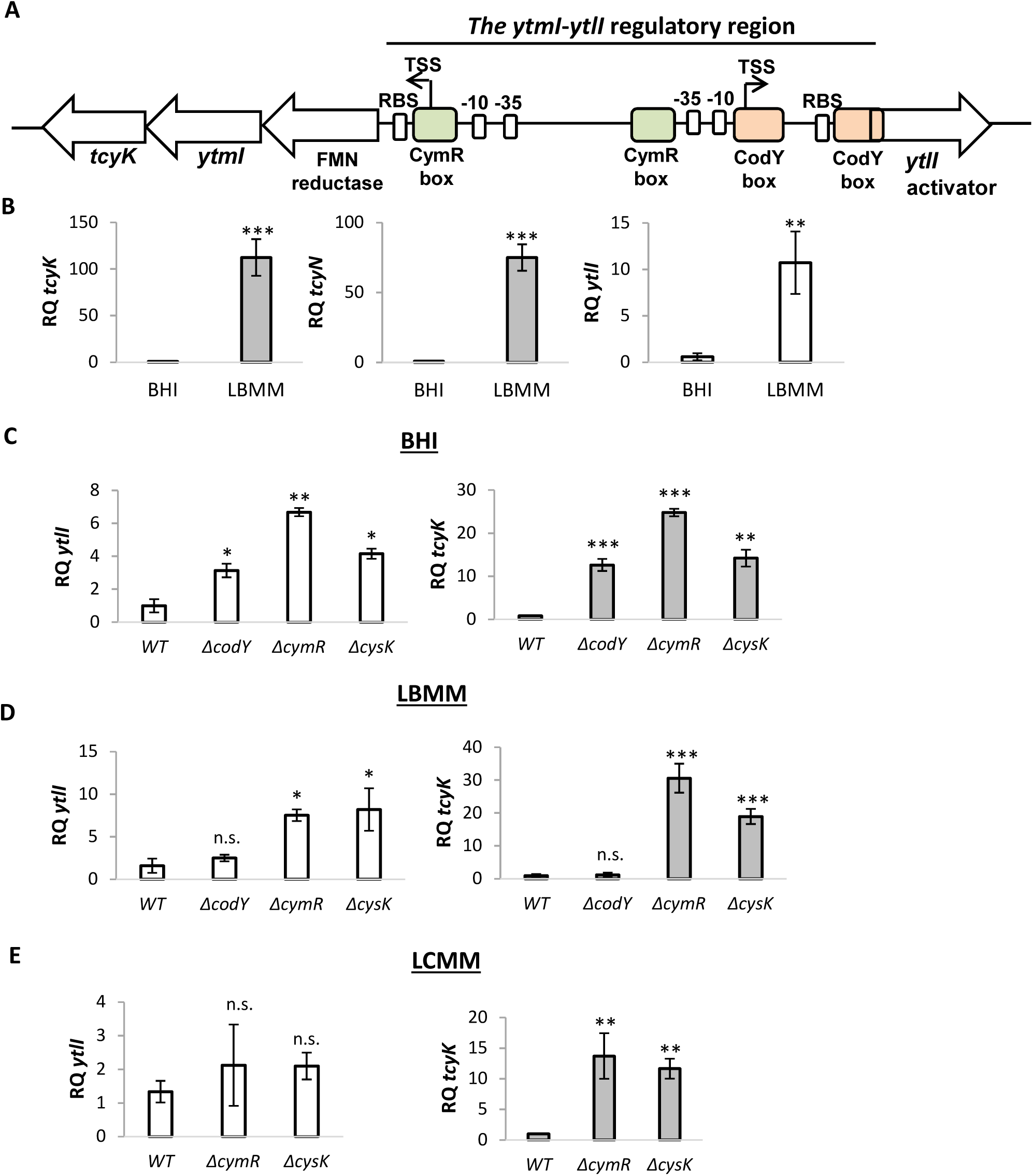
The TcyKLMN transporter is negatively regulated by CodY, CysK and CymR. **(A)** A schematic representation of the *ytmI-ytlI* regulatory region. Putative CymR binding-sites were identified using the motif of *Bacillus.* Putative CodY binding sites were identified using the motif of gram-positive bacteria. Promoters (−10 and −35) were identified using BPROM. TSS denotes the transcription start sites, either identified by Wurtzel *et al.,* 2012, or predicted based on the promoter sequence. **(B)** qRT-PCR analysis of *tcyK*, *tcyN* and *ytlI* transcription level in WT bacteria grown in rich media (BHI) and in LBMM. mRNA levels were normalized to *rpoD* mRNA and are represented as relative quantity (RQ), relative to mRNA level in WT bacteria grown in BHI. The data represent 3 biological replicates. Error bars indicate standard deviation. Asterisks represent P-values (* = P<0.05, ** = P<0.01, *** = P<0.001, n.s. = non- significant) calculated by Student’s t-test. P-values represent a comparison to the BHI sample. **(C-E)** qRT-PCR analysis of *tcyK* and *ytlI* transcription level in the indicated mutants grown in rich media (BHI) and minimal defined media supplemented with low concentration of either branched chain amino acids (LBMM) or cysteine (LCMM). mRNA levels were normalized to *rpoD* mRNA and are represented as relative quantity (RQ), relative to mRNA level in WT bacteria grown in the indicated medium. The data represent 3 biological replicates. Error bars indicate standard deviation. Asterisks represent P-values (* = P<0.05, ** = P<0.01, *** = P<0.001, n.s. = non- significant) calculated by Student’s t-test. P-values represent a comparison to the respective WT sample, unless indicated otherwise.

In light of these observations, we next examined whether the increased expression of the virulence genes in *ΔcymR* and *ΔcysK* is due to the increased expression of TcyKLMN. For this purpose, we generated double deletion mutants, lacking either *cysK* or *cymR* and *tcyK* (*ΔcysK/ΔtcyK* and *ΔcymR/ΔtcyK*) and analyzed their *Phly*-*lux* luminescence profiles during growth in LBMM. Notably, both mutants failed to activate *Phly*-*lux* expression, similarly to *ΔtcyK*, indicating that TcyKLMN itself plays a role in the activation of the virulence genes under low BCAA (Figure 4 and Figure S3 F-G).

**Fig 4.**
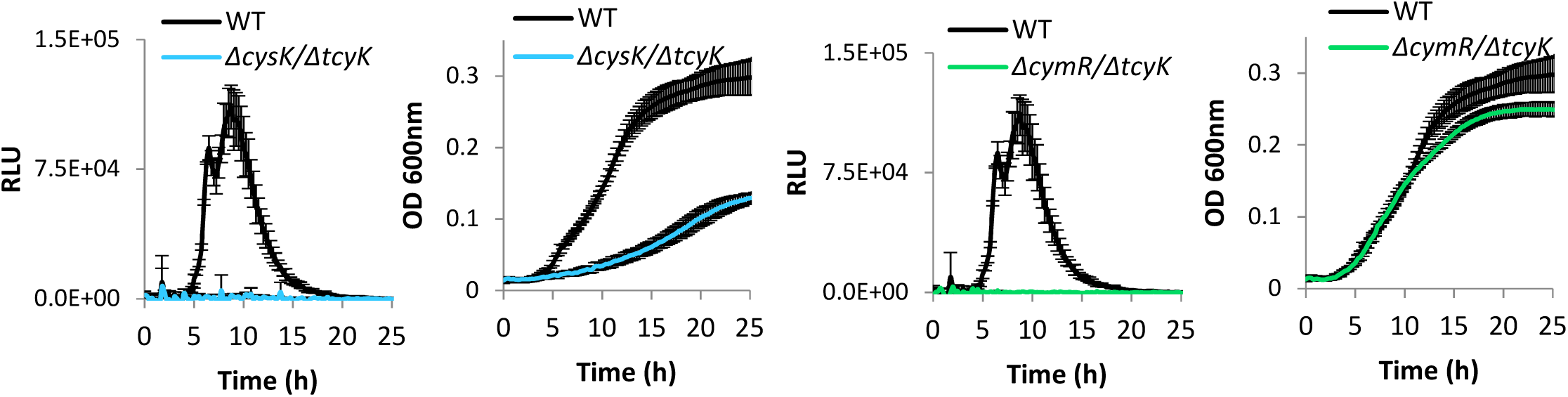
Enhanced expression of virulence genes in *ΔcymR* and *ΔcysK* is TcyKLMN dependent. Luminescence of pPL2 *Phly-lux* reporter system (left) and growth (right) of *ΔcysK/ΔtcyK* and *ΔcymR/ΔtcyK* bacteria grown in minimal defined media supplemented with low concentration of branched chain amino acids (LBMM). Relative Luminescence Units (RLU) represent luminescence values normalized to the respective OD 600nm value. OD 600nm values represent growth. The data represent 3 biological replicates. Error bars indicate standard deviation.

### Cysteine import via TcyKLMN drives glutathione biosynthesis, which triggers the induction of the virulence genes

Having discovered that in LBMM, TcyKLMN is required for the induction of the virulence genes and not for bacterial growth (Figure 1E), we hypothesized that its role in cysteine import may directly feed into glutathione biosynthesis, which, in turn, stimulates the activity of PrfA. This would account for the failure of all the tested transporter mutants (*i.e., ΔtcyK, tcyN*::*Tn, ΔcymR/ΔtcyK* and *ΔcysK/ΔtcyK*) to express the virulence genes in LBMM. To test this hypothesis, we first constructed a mutant deleted of the glutathione synthase gene (*ΔgshF*) and examined its impact on the activation of virulence genes in LBMM using the *Phly-lux* reporter system. As shown in Figure 5A, *ΔgshF* displayed practically zero expression of the *lux* genes (similarly to *ΔtcyK*), however this phenotype was fully restored to WT levels by the exogenous addition of GSH (20 mM). These observations indicated that glutathione biosynthesis is absolutely required for the activation of the virulence genes in LBMM. Similarly, the transcription of virulence genes in *ΔtcyK* was restored by the addition of GSH; *ΔtcyK* bacteria supplemented with 20mM of GSH exhibited a luminescence profile that was similar to that of WT bacteria, supporting the premise that TcyKLMN is involved in glutathione biosynthesis (Figure 5B). To directly examine this hypothesis, we measured the total concentration of glutathione (including reduced and oxidized forms, GSH + GSSG) in *ΔtcyK* bacteria in comparison to WT *Lm,* using *ΔgshF* as a control. Remarkably, the internal glutathione concentration of *ΔtcyK* was as low as that measured in *ΔgshF,* below the detection level of the glutathione assay kit (Figure 5C), indicating that TcyKLMN plays a role in glutathione biosynthesis, most likely by supplying the rate-limiting precursor, cysteine.

**Fig 5.**
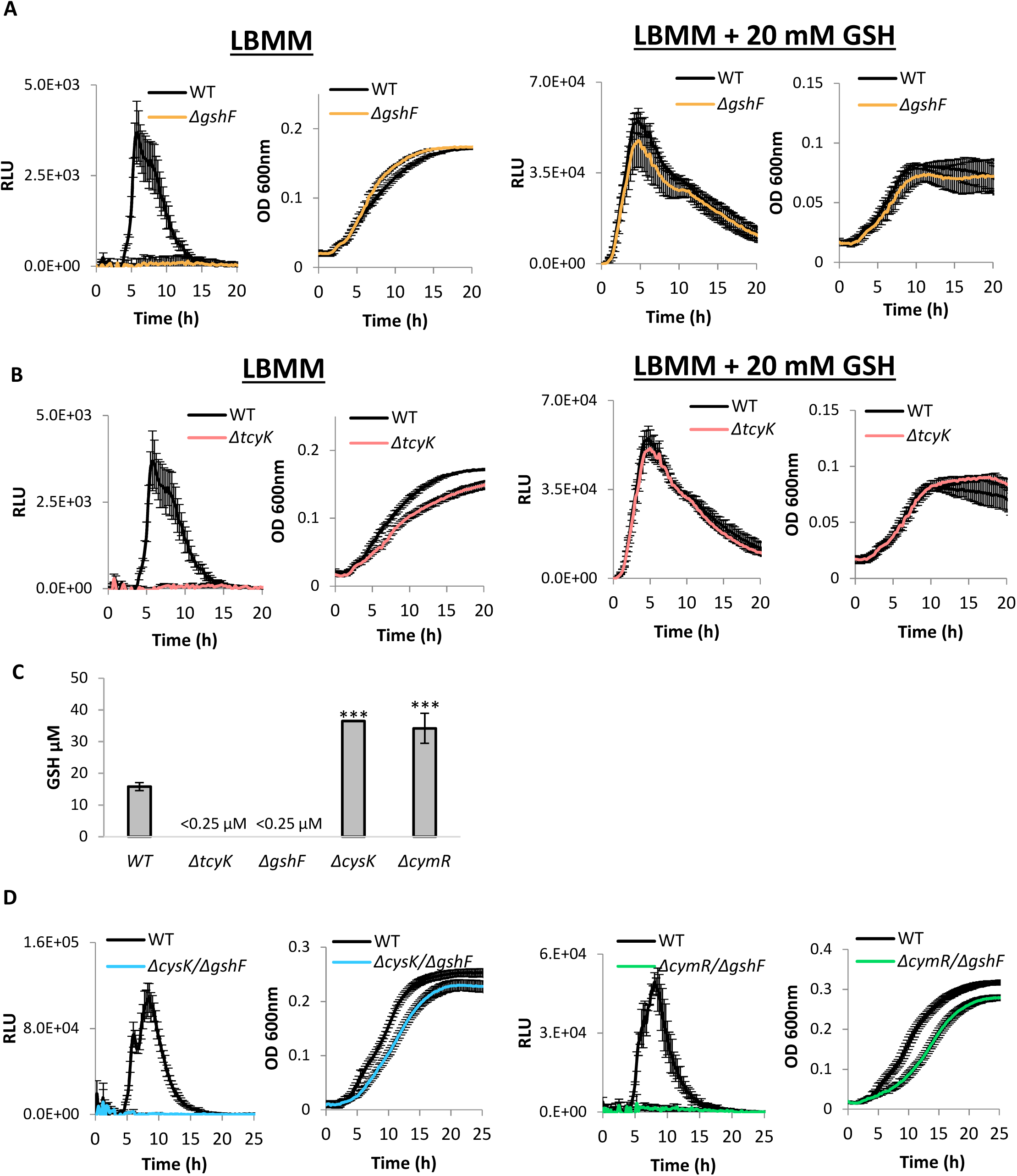
The TcyKLMN transporter plays a role in glutathione biosynthesis. **(A-B)** Luminescence of pPL2 *Phly-lux* reporter system (left) and growth (right) of *ΔgshF* and *ΔtcyK* bacteria grown in minimal defined media with low concentration of branched chain amino acids (LBMM). 20 mM reduced glutathione was supplemented to LBMM when indicated. Relative Luminescence Units (RLU) represent luminescence values normalized to the respective OD 600nm value. OD 600nm values represent growth. The data represent 3 biological replicates. Error bars indicate standard deviation. **(C)** Internal glutathione level during growth in LBMM in indicated bacteria. The data represent 3 biological replicates. Error bars indicate standard deviation. Asterisks represent P-values (*** = P<0.001) calculated by Student’s t-test. P-values represent a comparison to the WT sample. **(D)** Luminescence of pPL2 *Phly-lux* reporter system (left) and growth (right) of *ΔcysK/ΔgshF* and *ΔcymR/ΔgshF* bacteria grown in LBMM. Relative Luminescence Units (RLU) represent luminescence values normalized to the respective OD 600nm value. OD 600nm values represent growth. The data represent 3 biological replicates. Error bars indicate standard deviation.

In light of these findings, we next examined whether the enhanced expression of the virulence genes in *ΔcysK* and *ΔcymR* is a result of increased production of glutathione, as these strains highly express TcyKLMN, and hence presumably import more cysteine for glutathione synthesis. Notably, internal measurements of glutathione in *ΔcysK* and *ΔcymR* bacteria demonstrated a 2-fold increase in the amount of glutathione in comparison to WT bacteria, supporting the hypothesis that higher cysteine uptake can increase glutathione biosynthesis (Figure 5C). Moreover, these observations demonstrated that cysteine import by TcyKLMN feeds into glutathione biosynthesis and further confirmed that cysteine is indeed a rate-limiting substrate of GshF. To evaluate whether the enhanced virulence gene expression in *ΔcysK* and *ΔcymR* relies on glutathione biosynthesis, we combined these mutants with a deletion of *gshF* (generating the double deletion mutants: *ΔcysK/ΔgshF* and *ΔcymR*/*ΔgshF*), and examined virulence gene expression in LBMM, using the *lux* reporter system. As shown in Figure 5D, these mutants failed to express the *lux* genes, confirming that glutathione biosynthesis is absolutely required for virulence gene expression. Of note, these mutants, as well as *ΔgshF,* grew like WT bacteria both in BHI and LBMM, overall demonstrating that glutathione itself is not essential for growth under these conditions (Figure 5A, D and Figure S3 H-J). Finally, we examined whether CymR, CysK and CodY regulate the transcription of *gshF* in BHI and LBMM, and found that they do not (Figure S6), supporting the conclusion that they affect glutathione biosynthesis by regulating the import of cysteine via TcyKLMN. We also examined the possibility that TcyKLMN imports glutathione, and found that it does not, as evident from ITC experiments using TcyK, and growth experiments using glutathione as a source of cysteine (Figure S7 A-B). Of note, these experiments clearly demonstrated that *Lm* can use exogenous GSH as a sole source of cysteine (Figure S7B). Altogether, these findings demonstrated that when grown in LBMM, TcyKLMN is the main transport system that supplies cysteine for glutathione biosynthesis, which, in turn, drives the expression of the virulence genes via the activation of PrfA.

### TcyKLMN contributes to the induction of virulence genes during infection of macrophage cells

Finally, given the identified role of TcyKLMN in glutathione biosynthesis, we sought to investigate its impact on *Lm* intracellular growth and virulence gene expression in macrophage cells. We first analyzed the transcription level of *tcyK*, *tcyN* and *ytlI* in WT bacteria grown intracellularly in macrophage cells in comparison to bacteria grown in BHI. The data indicated a modest induction of TcyKLMN and its activator YtlI within the intracellular niche (∼3-fold) (Figure 6A). We next evaluated the expression of the virulence gene *plcA* in *ΔtcyK* and WT bacteria grown intracellularly in macrophage cells, using *ΔgshF* as a control. For this purpose, we used a fluorescence-based reporter system that we previously constructed, which relies on the expression of three consecutive *yfp* genes under the regulation of the *plcA* promoter (cloned in the integrative plasmid pPL2, pPL2-P*plcA*-3*yfp*) (33). Using this system, we observed a ∼40% reduction in the expression of *plcA* in *ΔtcyK* in comparison to WT bacteria, whereas a more dramatic reduction of ∼ 75% was observed in *ΔgshF* (Figure 6B). While these findings demonstrate that TcyKLMN contributes to the induction of the virulence genes during macrophage cell infection, they implied that it is not the sole supplier of cysteine within the intracellular niche, corroborating previous reports demonstrating that *Lm* also utilizes oligopeptides as a source of amino acids, including cysteine, which are imported by the Opp transporter (20, 41). We next examined the intracellular growth of *ΔtcyK* and *ΔgshF* in macrophage cells in comparison to WT bacteria. Notably, the data demonstrated a late growth defect for *ΔtcyK* that was apparent at 6 h post-infection (Figure 6C). Interestingly, a similar phenotype was reported for the Opp transporter deletion mutant (*ΔoppDF*), which also exhibited an intracellular growth defect at 6 to 7 h post-infection of macrophage cells (20). Evaluating *ΔgshF* intracellular growth in macrophage cells, we observed a more profound growth defect, starting at 3 h post-infection, indicating that glutathione biosynthesis by *Lm* is critical for intracellular growth, and hence has to be accompanied by cysteine uptake (Figure 6D). Taken together, the results presented here support the premise that *Lm* uses multiple transport systems to acquire cysteine in the intracellular niche, among them TcyKLMN.

**Fig 6.**
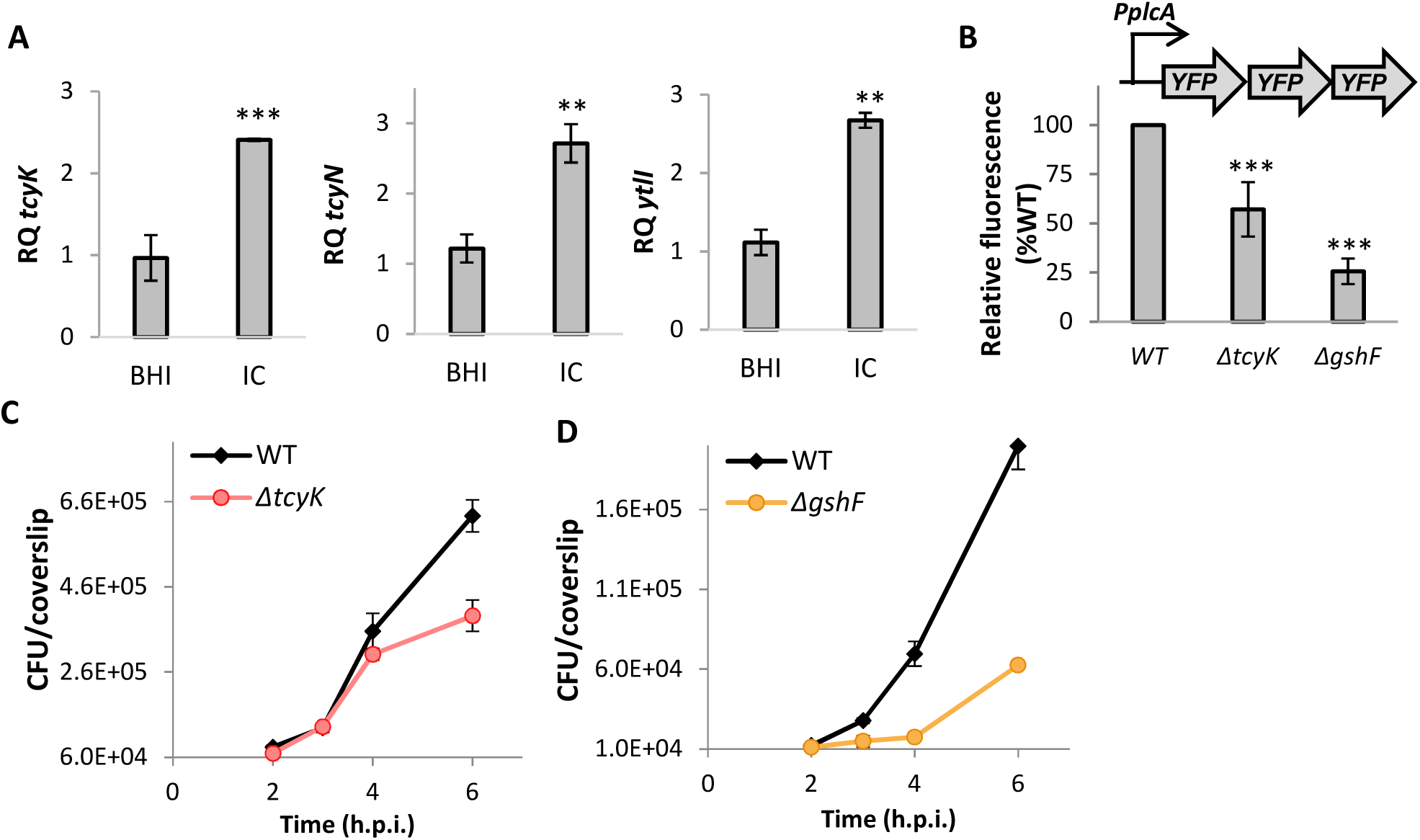
The TcyKLMN transporter promotes *Lm* intracellular growth in macrophage cells. **(A)** qRT-PCR analysis of *tcyK*, *tcyN* and *ytlI* transcription level in WT bacteria grown in rich media (BHI) or intracellularly in J774A.1 macrophage cell line (IC). mRNA levels were normalized to *rpoD* mRNA and are represented as relative quantity (RQ), relative to mRNA level in in the BHI sample. The data represent 3 biological replicates. Error bars indicate standard deviation. Asterisks represent P-values (* = P<0.05, ** = P<0.01, *** = P<0.001, n.s. = non- significant) calculated by Student’s t-test. P-values represent a comparison to the BHI sample. **(B)** Intracellular expression of *plcA* as indicated by the fluorescence of YFP proteins expressed under the control of *plcA* promoter (pPL2-*PplcA-3yfp*) in WT *Lm* and indicated mutants, grown in bone marrow derived macrophages (BMDM). Fluorescence was measured 3 hours post infection using a florescence microscope. The data represent at least 3 biological replicates. Error bars indicate standard deviation. Asterisks represent P-values (* = P<0.05, ** = P<0.01, *** = P<0.001, n.s. = non- significant) calculated by Student’s t-test. P-values represent a comparison to the WT sample. **(C-D)** Intracellular growth of *ΔtcyK* and *ΔgshF* mutants in BMDM cells in comparison to WT *Lm*. The experiment was repeated 3 times independently. Since we could not average the data, a representative result is shown here, and two additional biological repeats are shown in Figure S8. Error bars indicate standard deviation. h.p.i. = hours post infection.

## Discussion

This study is a direct continuation of our previous reports demonstrating that low availability of BCAA is sensed by *Lm* as a signal to activate the expression of virulence genes. While we established that the metabolic regulator CodY is responsible for the sensing of this signal, we further determined that it directly enhances the transcription of PrfA, thereby promoting virulence gene expression. That said, at the time these studies were conducted, we did not know that glutathione is also involved, through its binding to PrfA and allosterically enhancing its activity (28, 42). Here, we show that the induction of the virulence genes under low BCAA conditions is entirely dependent on *Lm*’s *de novo* synthesis of glutathione, mediated by GshF. Further, we demonstrate that glutathione biosynthesis under this condition requires the active import of cysteine. As indicated, *Lm* is auxotrophic to cysteine and methionine and hence has to import these amino acids from the environment. To date, two transport systems have been linked to the acquisition of cysteine in *Lm* (as a free amino acid or in peptides), yet their role in supporting *Lm* intracellular growth was not robust, implying that other systems may be involved. Interestingly, studying the regulation of *Lm* virulence gene expression under low BCAA (i.e., in LBMM) led to the discovery of TcyKLMN as a cystine/cysteine importer that facilitates glutathione biosynthesis and PrfA activation. The results demonstrated that TcyKLMN is repressed under nutrient-rich conditions, and up-regulated when BCAA are limited. This repression was linked to CodY, which we found to repress the activator of the *ytmI* operon, YtlI. As indicated, low BCAA is the same condition at which PrfA expression is up-regulated by CodY; therefore, it is possible that the parallel increase in cysteine uptake, and subsequent up-regulation of glutathione biosynthesis, may have evolved to cope with the heightened PrfA levels. Altogether, the findings presented here reveal an intriguing synchronization between PrfA expression and glutathione biosynthesis that is mediated by CodY, overall indicating that CodY regulation of *Lm* virulence is even more complex than previously considered.

An important observation made in this study was that glutathione levels are largely limited within the bacteria, and that the availability of cysteine determines its biosynthesis. In line with this conclusion, all the factors that were found to affect cysteine import, i.e., CodY, CymR, CysK and TcyKLMN, indirectly influenced the activation of PrfA. In a way, these findings uncover a new pathway by which *Lm* virulence can be regulated in the mammalian host responding to changes in cysteine availability. CodY, CymR and CysK were also found to regulate the expression of TcyKLMN, and hence to modulate the expression of the virulence genes. These regulators still repressed TcyKLMN under conditions of low concentrations of BCAA and cysteine, implying that other metabolic or environmental signals may be involved. In *B. subtilis*, it was shown that the *ytmI* operon is differentially regulated under disulfide stress and changes in sulfur availability, hence it is possible that the *Lm* CymR and CysK respond to these conditions (36, 38). Moreover, since glutathione plays a role in redox conditions, it is possible that CymR and CysK respond to these conditions as well. In this respect, it was demonstrated in *Staphylococcus aureus* that CymR senses oxidative stress via thiolation of its cysteine residue (Cys-25), and hence the YtmI operon is up-regulated under oxidizing environments (43). Taken together, these reports indicate that the YtmI operon is regulated by multiple metabolic and environmental cues. Interestingly, the *ytmI* operon of *B. subtilis* is independent of CodY regulation, as evident from RNA-seq experiments and genome-wide analyses of CodY binding sites (44, 45). It is tempting to speculate that CodY regulation of *Lm*’s TcyKLMN evolved to cope with the pathogenic lifestyle of this bacterium, yet *Lm* and *B. subtilis* differ greatly in their sulfur, cysteine and methionine metabolism, which can lead to differences in the regulation of these pathways. Notwithstanding, this study revealed an intriguing link between CodY regulation, cysteine import and glutathione biosynthesis in *Lm*, portraying additional mechanism to control the expression of the virulence genes.

Here, we showed that the SBP of TcyKLMN (i.e., TcyK) specifically binds cysteine, and to a better extent, CSSC, suggesting that it primes the import of these two related substrates. CSSC is the oxidized form of cysteine, containing two cysteine molecules that are linked via a disulfide bond, and hence is expected to be more prevalent in oxidizing environments, such as within phagosomes. Outside the mammalian cells (i.e., in the extracellular milieu), cysteine is considered to be abundant (produced and secreted by the liver), yet it is quickly oxidized and imported into the cells via specialized CSSC transporters, e.g. the CSSC/glutamate antiporter, cXT, which also plays a role in glutathione synthesis (46–49). Interestingly, within the cells, thioredoxin or glutathione are used to reduce the CSSC to cysteine, which is further utilized in protein synthesis. Of note, mammalian cells can further biosynthesize cysteine from methionine, using the transsulfuration pathway, or alternatively break down glutathione to salvage cysteine (46). In respect to *Lm*’s intracellular lifestyle, it is likely that it encounters CSSC within the phagosome/vacuole environment. It is possible that TcyKLMN plays a role during cell invasion or during spread from cell to cell (i.e., in the primary and secondary vacuoles, respectively), or when *Lm* switches to persistent infection, i.e., when it resides in lysosome-like vacuoles for long periods (50, 51). While we did not identify a phenotype for *ΔtcyK* upon mice infections (data not shown), we observed a late intracellular growth defect during infection of bone marrow-derived macrophage cells. These experiments further demonstrated a reduced transcription level of *plcA* in *ΔtcyK*, supporting the premise that TcyKLMN contributes to the activation of PrfA during *Lm* infection of mammalian cells. Since CSSC and cysteine are relatively limited in the intracellular environment, it is not surprising that *Lm* acquired multiple means to scavenge this essential amino acid. As mentioned, *Lm* exploits peptides as a source of amino acids, and the Opp transporter was shown to import cysteine-containing peptides as a source of cysteine for glutathione biosynthesis (20, 41). The finding that *ΔtcyK* exhibits a 40% reduction in *plcA* transcription, as compared to the 75% reduction in *ΔgshF*, supports the premise that *Lm* exploits multiple systems to scavenge cysteine.

Reniere et al. suggested that *Lm* also exploits host glutathione (GSH) to activate PrfA (17). However, this mode of PrfA activation appeared to be minor, as *ΔgshF* bacteria hardly activated the virulence genes in the intracellular environment (17), overall indicating that host-derived GSH cannot replace the glutathione produced by *Lm*. In this regard, it remains an open question why *Lm* synthesizes GSH and uses it as an intracellular signal, considering its high abundance in the intracellular niche, much more than cysteine. Other bacterial pathogens, such as *Burkholderia pseudomallei* and *Francisella tularensis* were shown to exploit host GSH as an activating signal of virulence gene expression and as a source of cysteine, respectively (52, 53). Intriguingly, *in vitro* experiments in minimal defined medium indicated that *Lm* is capable of importing GSH and uses it both to activate PrfA, and as a source of cysteine (Figure 5 **and** S7B) (54). While these findings imply that *Lm* encodes a glutathione transporter, or, alternatively, uses a nonspecific system such as di/tri-peptide transporters to import glutathione, it is not clear why it does not use them in the intracellular niche. Despite its importance, a dedicated glutathione transporter was not identified in Gram-positive bacteria. In Gram-negative bacteria the dipeptide transporter DppBCDF was shown to import glutathione, using the SBP protein GbpA (55). It is of particular interest to decipher the structural and molecular mechanism by which glutathione is imported in Gram-positive bacteria, especially intracellular pathogens like *Lm*. Intriguingly, in the Gram-positive bacterium *Streptococcus mutans*, it was shown that the CSSC ABC-transporter, TcyBC (not found in *Lm*), imports both CSSC and glutathione using two distinct SPB proteins (55). The SBP that binds glutathione (GshT) was found to be encoded elsewhere on the bacterial chromosome, and not near the *tcyBC* genes. While this phenomenon of a shared permease is not new, the discovery that *S. mutans* holds a distinct SBP that primes glutathione import via another transporter was new, providing early insights into glutathione import in Gram-positive bacteria (56). Interestingly, we found that TcyK shares 30% amino-acid sequence identity with the *S. mutans* GshT, yet our data indicated that it does not bind glutathione.

In summary, this study demonstrated that cysteine import is critical for virulence gene expression in *Lm*. The data imply that multiple metabolic and environmental cues are involved in the regulation of cysteine import and hence affect glutathione biosynthesis, placing this function at the heart of *Lm* patho-metabolism. As different bacterial pathogens acquired different mechanisms to sense the mammalian niche, it is interesting to learn how they exploit host and bacterial metabolites as signals and effectors of virulence.

## Materials and Methods

### Ethics statement

Experimental protocols were approved by the Tel Aviv University Animal Care and Use Committee (01-15-052, 04-13-039) and were in accordance with the Israel Welfare Law (1994) and the National Research Council guide (Guide for the Care and Use of Laboratory Animals 2010).

### Bacterial strains, plasmids and primers

*Listeria monocytogenes* (*Lm*) strain 10403S was used as the WT strain and as the parental strain to generate allelic exchange mutant strains (**Table S1**). *E. coli* XL-1 blue strain (Stratagene) was used to generate vectors and *E. coli* SM-10 strain (57) was used for plasmid conjugation to *Lm.* Plasmids and primers used in this study are listed in **Table S1** and **Table S2**, respectively.

### Growth conditions

*Lm* bacteria were grown at 37°C, with agitation, in brain heart infusion (BHI), as a nutrient-rich medium, or in minimal defined medium (MM). MM (phosphate buffer 48.2 mM KH_2_PO_4_ and 1.12 M Na_2_HPO_4_, pH 7, 0.41 mg/ml magnesium sulfate, 10 mg/ml glucose, 100 μg/ml of each amino acid (methionine, arginine, histidine, tryptophan, phenylalanine, cysteine, isoleucine, leucine and valine), 600 µg/ml glutamine, 0.5 mg/ml biotin, 0.5 mg/ml riboflavin, 20 mg/ml ferric citrate, 1 mg/ml para-aminobenzoic acid, 5 ng/ml lipoic acid, 2.5 mg/ml adenine, 1 mg/ml thiamine, 1 mg/ml pyridoxal, 1 mg/ml calcium pantothenate and 1 mg/ml nicotinamine) was prepared as described previously (58). For analysis of auxotrophies, *Lm* was grown with 0-2000 μg/ml of either cysteine or methionine or neither, as indicated. For growth under low BCAA, MM was freshly made with 10-fold less isoleucine, leucine and valine (resulting in a final concentration of 10 μg/ml) was named low-BCAA minimal defined medium (LBMM). For growth under limited concentrations of cysteine, MM was freshly made with 10-fold less cysteine (resulting in a final concentration of 10 μg/ml). For growth in glutathione medium, 20 mM reduced glutathione (GSH) were freshly added to LBMM. For growth with cysteine or with either cystine (CSSC), reduced or oxidized glutathione (GSH or GSSG, respectively) as a cysteine source, MM was prepared without cysteine and 0-2 mM of the respective cysteine source was added to the medium.

### Lux reporter assay

Overnight BHI cultures harboring a *Phly*-*lux* luciferase reporter system (pPL2*-Phly-luxABCDE*) (59) were washed 3 times with PBS, adjusted to OD_600_ = 0.03 in fresh LBMM or GSH medium, and grown in a 96-well plate. Luminescence and bacterial growth (OD_600_) were measured every 15 min after shaking for 1 min, using a Synergy HT BioTek plate reader at 37°C for 12-48 h.

### Bacterial RNA extraction and qRT-PCR

Bacteria grown in the indicated medium were harvested at mid-logarithmic growth (OD_600_ of ∼0.3). Total RNA was extracted from bacteria using the RNA*snap* method (60). Briefly, bacterial pellets were washed with AE buffer (50 mM NaOAc pH 5.2, 10 mM EDTA) and then resuspended in 95% formamide, 18 mM EDTA, 1% 2-mercaptoethanol and 0.025% SDS. Bacterial lysis was performed by vortexing extracts with 100 µm zirconia beads (OPS Diagnostics), followed by incubation at 95°C. Nucleic acids were precipitated with ethanol and treated with Turbo-DNase (Ambion), followed by standard phenol extraction. Total RNA (1 µg) was reverse-transcribed to cDNA using qScript (Quanta). qRT-PCR was performed on 10 ng cDNA using FastStart Universal SYBR Green Master (Roche) in a StepOne Plus real-time PCR system (Applied Biosystems). The transcription level of each gene was normalized to that of the reference gene *rpoD*.

### Analysis of promoters and putative binding sites

Putative binding sites of transcriptional regulators were identified using the MAST search tool (61), using the CymR motif of *Bacillus* (38) and CodY motif of Gram-positive bacteria (62). Promoters (−10 and −35) were identified using BPROM (63). Transcription start sites (TSS) were either previously identified (64) or manually predicted based on the promoter sequence.

### Intracellular growth in macrophage cells

Bone marrow-derived macrophages (BMDM) were used for *Lm* infection experiments. The cells were isolated from 8 week-old female C57BL/6 mice (Envigo, Israel) and cultured in BMDM medium (Dulbecco’s modified Eagle medium (DMEM) supplemented with 20% fetal bovine serum, sodium pyruvate (1 mM), L-glutamine (2 mM), β-mercaptoethanol (0.05 mM) and monocyte-colony stimulating factor (M-CSF, L929-conditioned medium), as described previously (65)). BMDM cells (2×10^6^) were seeded in a 60 mm Petri dish, on glass coverslips, in 5 ml BMDM medium and incubated overnight (O.N.) in a 37 °C, 5% CO_2_ forced-air incubator. *Lm* bacteria (8×10^6^) grown O.N. at 30 °C without agitation, were used to infect BMDM cells (MOI of 1). Thirty minutes post-infection, macrophage monolayers were washed and fresh medium was added. Gentamicin was added 1 h post-infection to a final concentration of 5 μg/ml in order to limit the growth of extracellular bacteria. At each time point, three coverslips were transferred to 2 ml sterile water to release the intracellular bacteria. Serial dilutions of the resulting lysate were plated on BHI agar plates and the CFUs were counted after 24 h incubation at 37 °C.

### Intracellular P*plcA*-3*yfp* expression

WT and mutant strains expressing three consecutive YFP proteins under the regulation of the *plcA* gene promoter (cloned on the pPL2 integrative plasmid) were used to infect BMDM on 20 mm slides. Three hours post-infection, cells were fixed with 4% v/v paraformaldehyde-PBS and permeabilized with 0.05% v/v Triton X-100. DNA was stained with DAPI-containing Vectashield mounting medium (Vector laboratories inc.). Fluorescent images were captured using a Zeiss LSM 510-META confocal microscope.

### Intracellular gene expression analysis

RNA was purified from WT *Lm* bacteria intracellularly grown in J774A.1 macrophage cells, as previously described for BMDM macrophages (66). Briefly, three 145 mm dishes were seeded with 2×10^7^ cells that were then infected in parallel with 2×10^9^ bacteria. Thirty minutes post-infection, cell monolayers were washed twice with PBS to remove unattached bacteria, and fresh medium was added. At 1 h post-infection (h.p.i.), gentamicin (50 μg/ml) was added to limit extracellular bacterial growth. At 6 h.p.i., the macrophages were lysed with 20 mL cold water, and cell debris and nuclei were removed by centrifugation (800 xg, 3 min, 4°C). Released bacteria were quickly collected on 0.45μm filter membranes (Millipore) using a vacuum apparatus and snap-frozen in liquid nitrogen. Bacteria were recovered from the filters by vortexing the membranes in AE buffer (50 mM sodium acetate pH 5.2, 10 mM EDTA), and bacterial nucleic acids were extracted using the *RNAsnap* method (60), followed by ethanol precipitation. RNeasy Mini Kit DNase on column (QIAGEN) was used for DNase treatment.

### Cloning, overexpression and purification of TcyK

*Lm* 10403S *tcyK*, encoding the TcyKLMN substrate binding protein, was synthesized and adjusted to the *E. coli* codon usage (Genescript). The gene was cloned into the pET-19b vector (Novagen) for expression with an N-terminal His-tag. His-tagged TcyK was overexpressed in *E. coli* BL21-Gold (DE3) (Stratagene) cultured in Terrific Broth (TB) and induced at mid-log phase by addition of 1 mM isopropyl b- D-1-thiogalactopyranoside (IPTG; 1.5 h, 37°C). Cells were harvested by centrifugation (13,600 × g, 20 min, 4°C) and the pellet was stored at -80°C until use. For purification, cells were homogenized in 50 mM Tris-HCl pH 8, 500 mM NaCl, complete EDTA-free protease inhibitor (Roche), 30 mg ml-1 DNase (Worthington), and 1 mM MgCl_2_. The cells were then ruptured by three passages through an EmulsiFlex-C3 homogenizer (Avestin), and the lysate was centrifuged at 34,500 × g for 30 min, at 4°C. The supernatant was loaded onto a nickel affinity column (HisTrap HP, GE Healthcare) on an AKTA Avant instrument. The protein was eluted using an imidazole gradient, and imidazole was removed from the sample by desalting (HiPrep 26/10, GE Healthcare). Protein purification was monitored by Coomassie staining of SDS-PAGE and size exclusion chromatography (Superdex 75 10/300 GL, GE Healthcare).

### Isothermal titration calorimetry experiments

Calorimetric measurements were performed with the Microcal iTC200 System (GE Healthcare). Prior to measurements, purified TcyK was dialyzed against three exchanges of 50 mM Tris-HCl pH 8, 500 mM NaCl buffer. Stocks of cysteine, CSSC, GSSG, GSH or control amino acid (Gln, His, Glu, Thr and Ser) were prepared fresh in double-distilled water and diluted to a working concentration using the buffer from the last protein dialysis exchange. 1 mM Tris(2-carboxyethyl)phosphine (TCEP) was added to cysteine and GSH stocks as a reductant agent. Aliquots (2 μL) of 500 μM stocks were added by a rotating syringe to the reaction well containing 200 μL of 50 μM purified TcyK at 25°C. Data-fitting was performed with the Microcal analysis software.

### Glutathione quantification

Total glutathione (GSH+GSSG) concentration in bacteria was measured as described previously (54). Briefly, bacteria were grown to mid-log phase in LBMM, resuspended in PBS containing 1 mM EDTA, pH 6.5, and lysed by vortexing with 100 µm zirconia beads (OPS Diagnostics). Cold (4°C) lysates were deproteinated with an equal volume of metaphosphoric acid and quantified with a commercial kit (Cayman Chemical), according to the recommendations of the kit manufacturer.

## Acknowledgments

We thank Lior Lobel for critical review of the manuscript. This work was supported by the NIH grant (R01AI109048) of the US National Institute of Allergy and Infectious Diseases, granted to Anat A Herskovits, Abraham L. Sonenshein and Mary O’Riordan.

---

**Fig S1.**
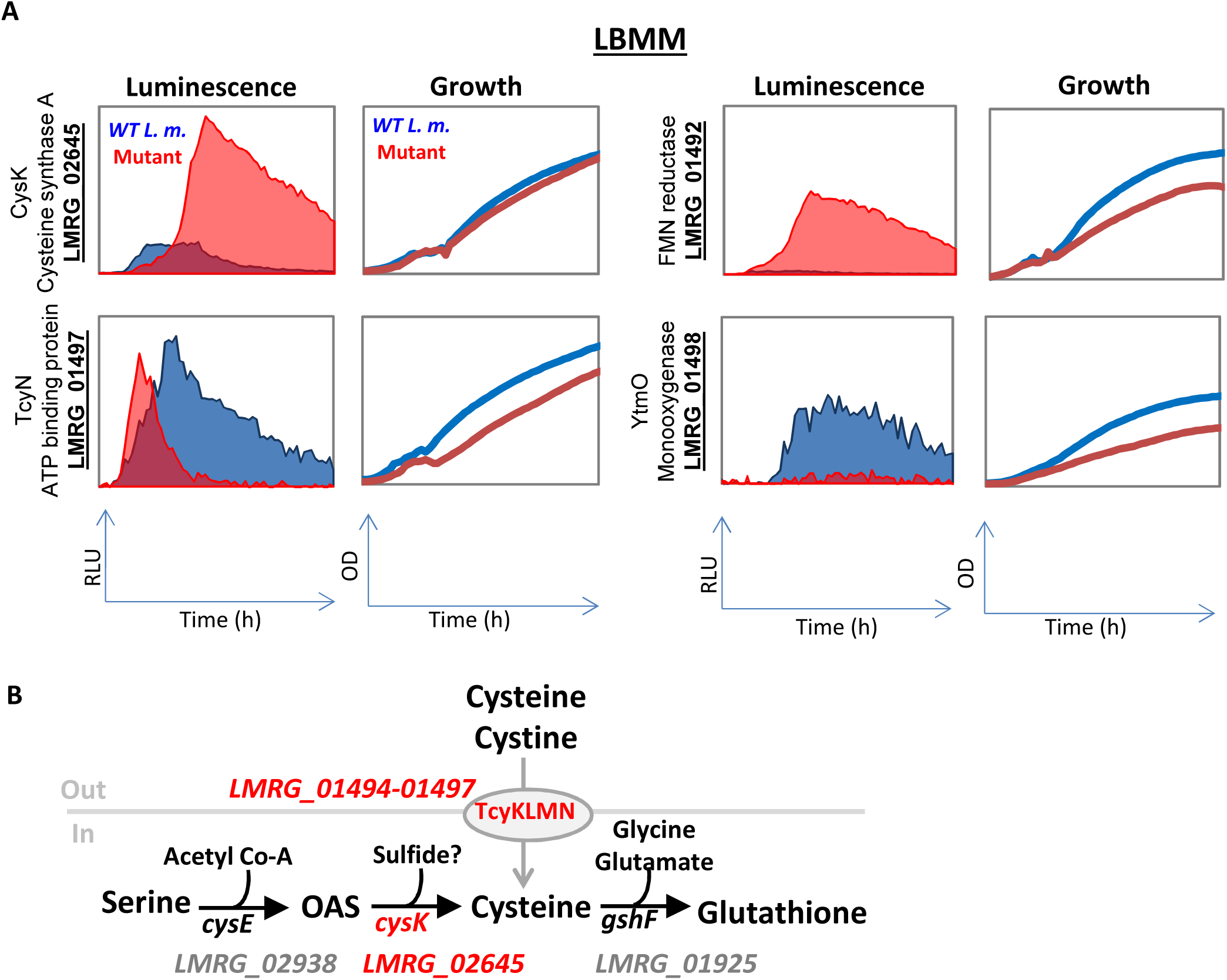
Cysteine uptake and metabolic genes identified in a screen by Friedman *et al*. **(A)** Phenotypes of transposon mutants as identified in Friedman *et al.,* 2017. WT bacteria and indicated transposon mutants were grown in minimal defined medium supplemented with low concentration of branched chain amino acids (LBMM). Relative Luminescence Units (RLU) represent luminescence values normalized to the respective OD 600nm value. **(B)** Schematic representation of cysteine transport via TcyKLMN and cysteine-glutathione biosynthesis. The genes identified in the genetic screen are marked in red. OAS=O-acetylserine.

**Fig S2.**
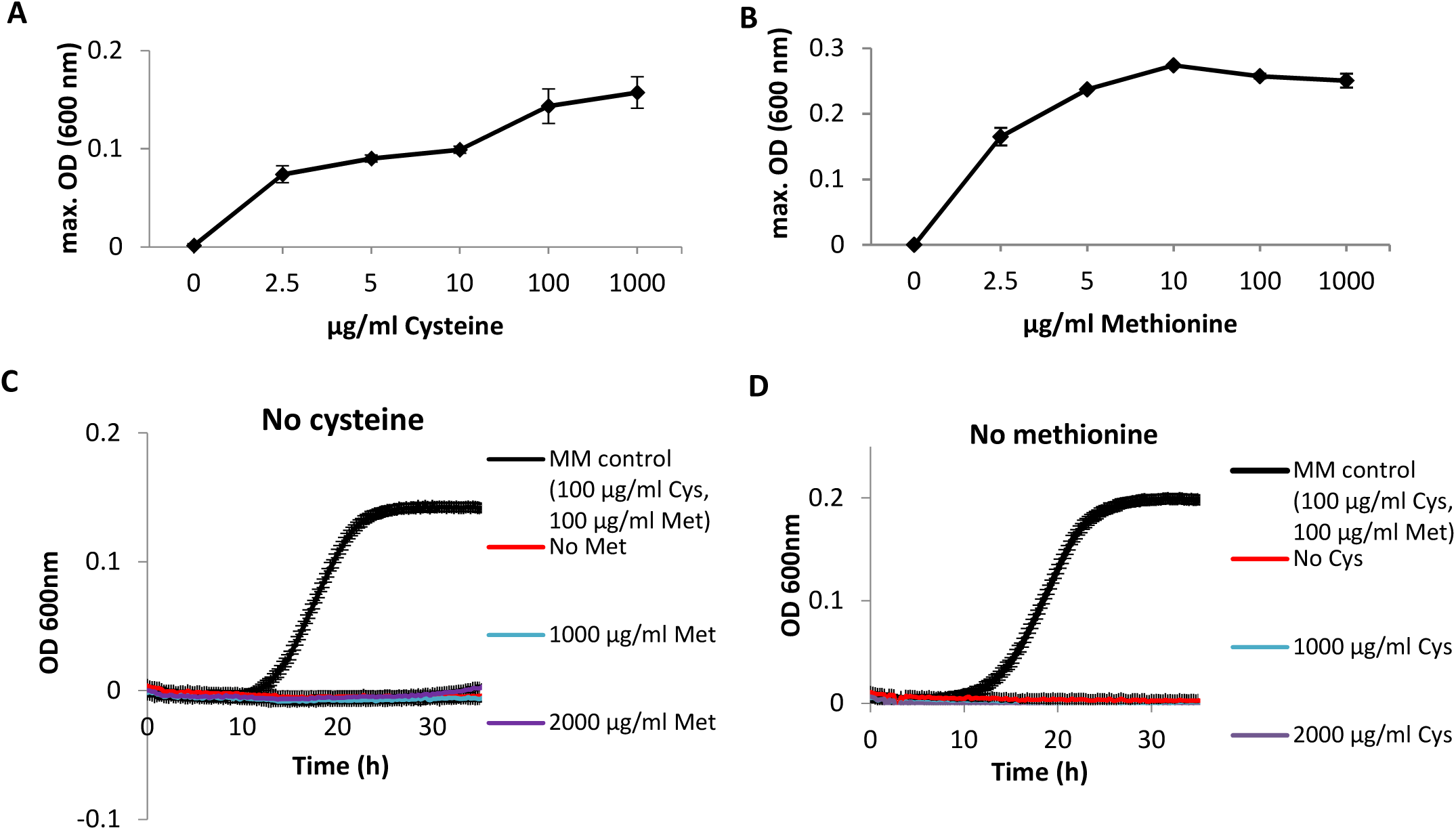
*Lm* strain 10403S is auxotrophic to cysteine and methionine. **(A-B)** Growth of WT *Lm* strain 10403S in minimal defined media supplemented with different concentrations of either cysteine or methionine. The data is presented as maximal OD 600nm values, corresponding to bacterial yield. The data represent 3 biological replicates. Error bars indicate standard deviation. **(C)** Growth of WT *Lm* in minimal defined media without cysteine supplemented either with 1000 µg/ml, 2000 µg/ml or no methionine. Growth of WT *Lm* in minimal defined media supplemented with standard concentrations of cysteine (100 µg/ml) is shown as a control. OD 600 nm values represent growth. The data represent 3 biological replicates. Error bars indicate standard deviation. **(D)** Growth of WT *Lm* in minimal defined medium without methionine, supplemented with either 1000 µg/ml, 2000 µg/ml or no cysteine. Growth of WT *Lm* in minimal defined media supplemented with standard concentrations of methionine (100 µg/ml) is shown as a control. OD 600nm values represent growth. The data represent 3 biological replicates. Error bars indicate standard deviation.

**Fig S3.**
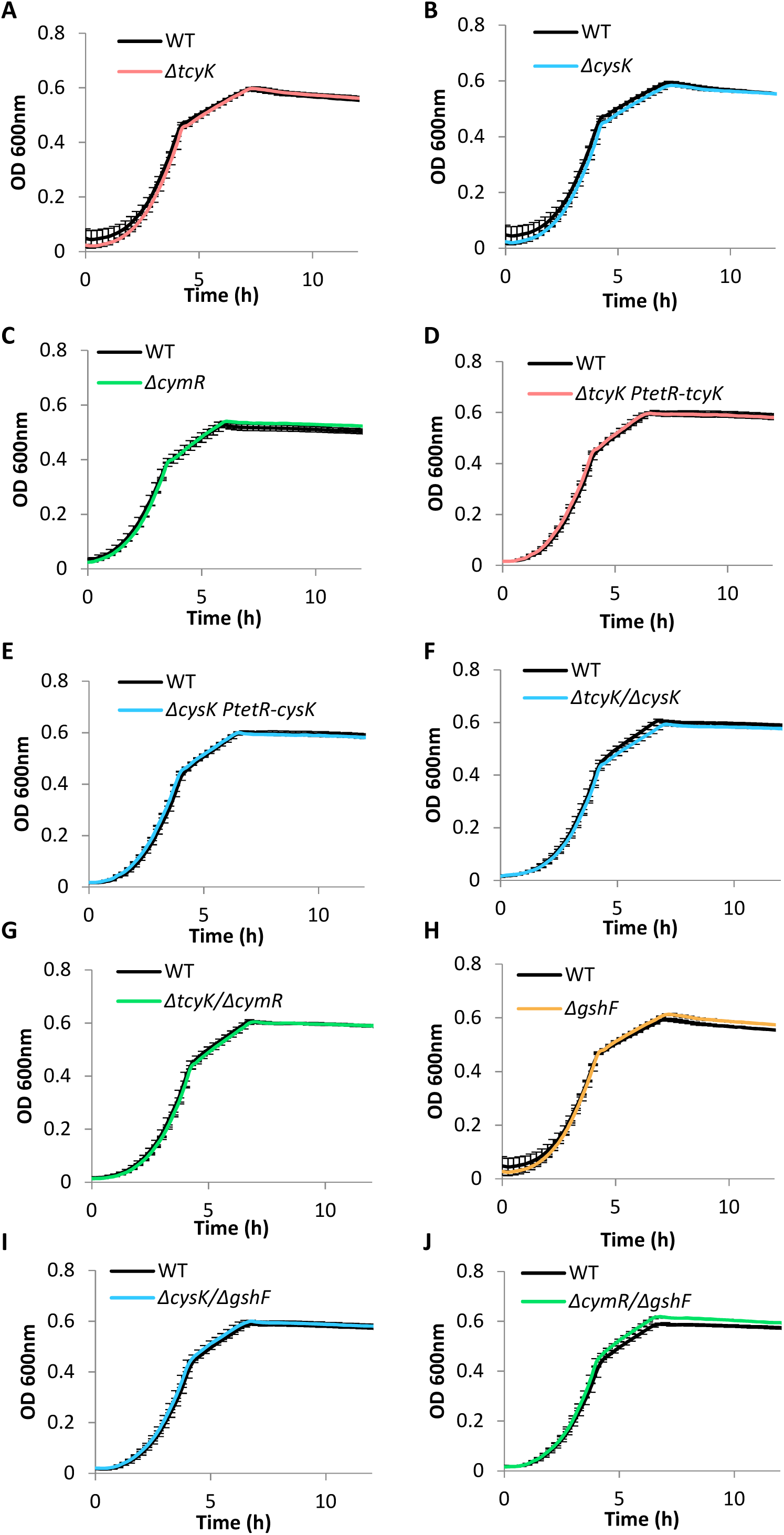
Growth of deletion mutants generated in this study in the rich medium BHI. WT bacteria and indicated strains **(A-J)** were grown in BHI. OD 600nm values represent growth. The data represent 3 biological replicates. Error bars indicate standard deviation.

**Fig S4.**
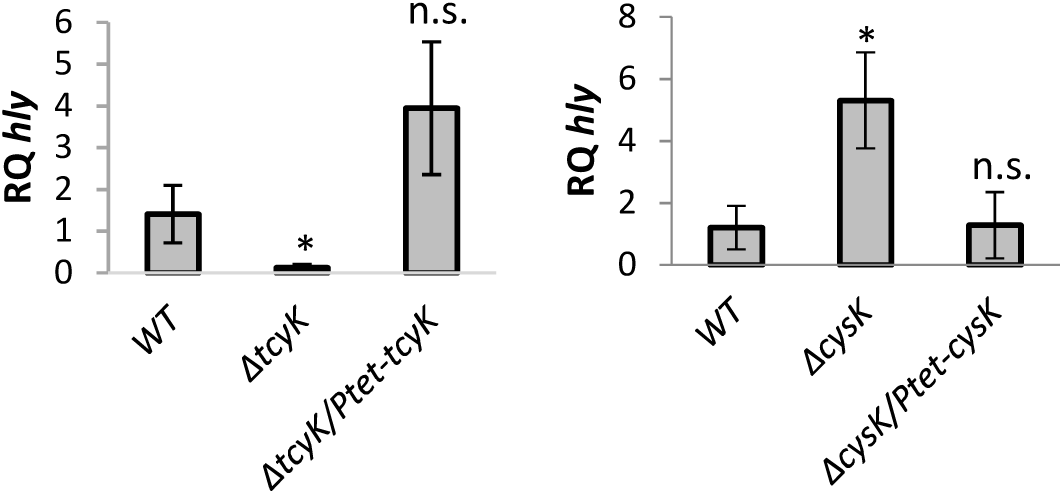
Complementation experiments of *ΔtcyK* and *ΔcysK* mutants. qRT-PCR analysis of *hly* transcription level in WT *Lm*, *ΔtcyK* and *ΔcysK* mutants and their respective complementation strains harboring a copy of *tcyK* or *cysK* gene under the *tet* promoter on pPL2 plasmid, grown in LBMM. mRNA levels were normalized to *rpoD* mRNA and are represented as relative quantity (RQ), relative to mRNA level in WT bacteria. The data represent at least 2 biological replicates. Error bars indicate standard deviation. Asterisks represent P-values (* = P<0.05, ** = P<0.01, *** = P<0.001, n.s. = non- significant) calculated by Student’s t-test. P-values represent a comparison to the WT sample unless indicated otherwise.

**Fig S5.**
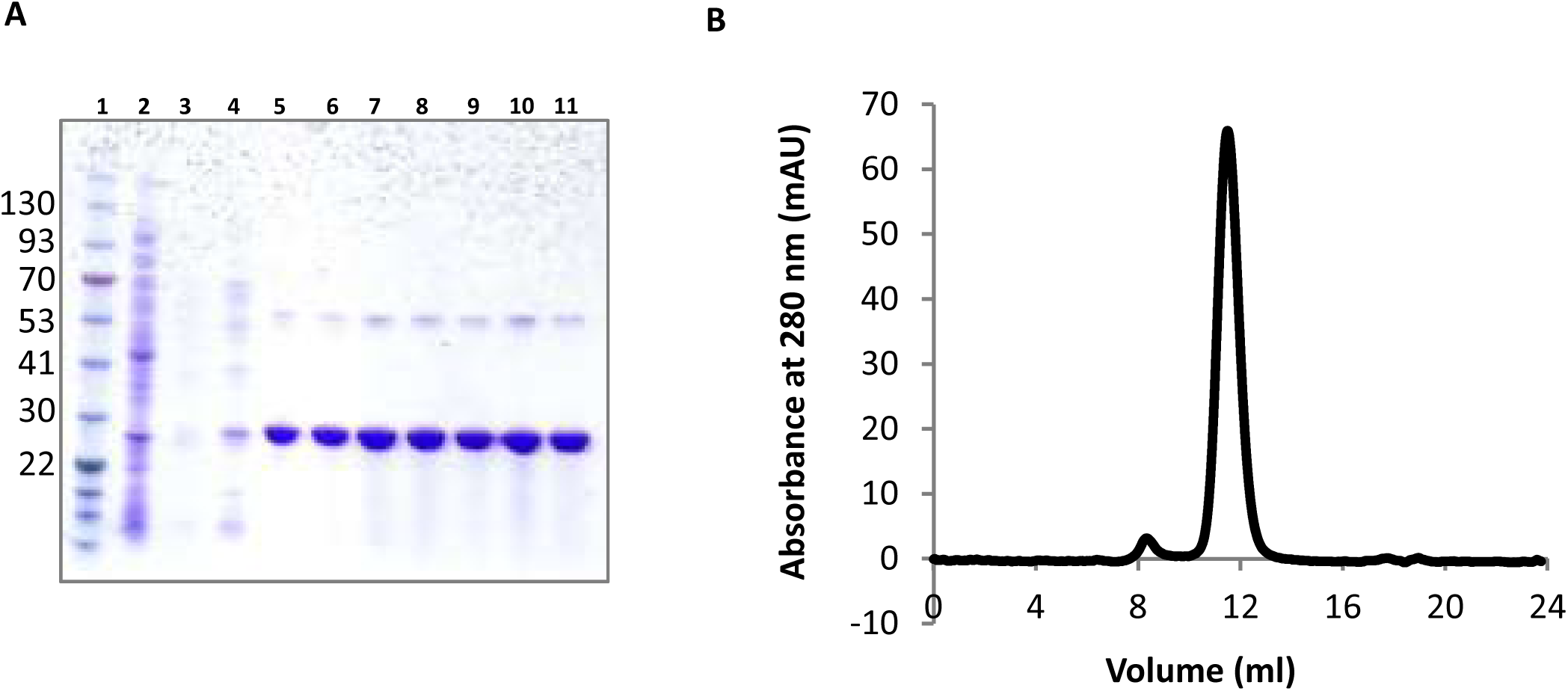
Purification of a short variant of TcyK. **(A)** Coomassie staining of SDS-PAGE of Ni-NTA affinity purification of a His-tagged short TcyK (amino acids 38-286). Lane 1: molecular weight marker (in kDa), Lane 2: total protein extract, Lane 3: column-unbound fraction, Lane 4: wash with 60 mM of imidazole, Lanes 5-11: fractions eluted using a linear gradient of 60 to 250 mM imidazole. **(B)** Analysis of the purified protein by size exclusion chromatography. 80 µl of a 0.8 mg/ml protein solution were injected on a Superdex 75 10/300 GL column (GE Healthcare).

**Fig S6.**
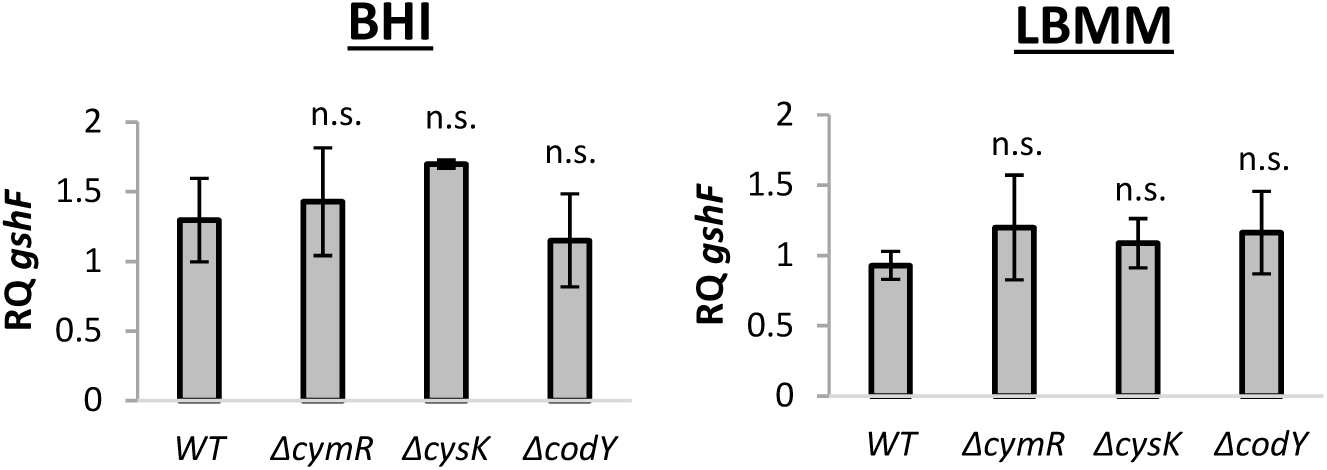
*gshF* is not transcriptionally regulated by CymR, CysK and CodY. qRT-PCR analysis of *gshF* transcription level in the indicated mutants grown in rich media (BHI) and minimal defined media supplemented with low concentration of either branched chain amino acids (LBMM). mRNA levels were normalized to *rpoD* mRNA and are represented as relative quantity (RQ), relative to mRNA level in WT bacteria grown in the indicated medium. The data represent at least 2 biological replicates. Error bars indicate standard deviation Asterisks represent P-values (* = P<0.05, ** = P<0.01, *** = P<0.001, n.s. = non- significant) calculated by Student’s t-test. P-values represent a comparison to the respective WT sample.

**Fig S7.**
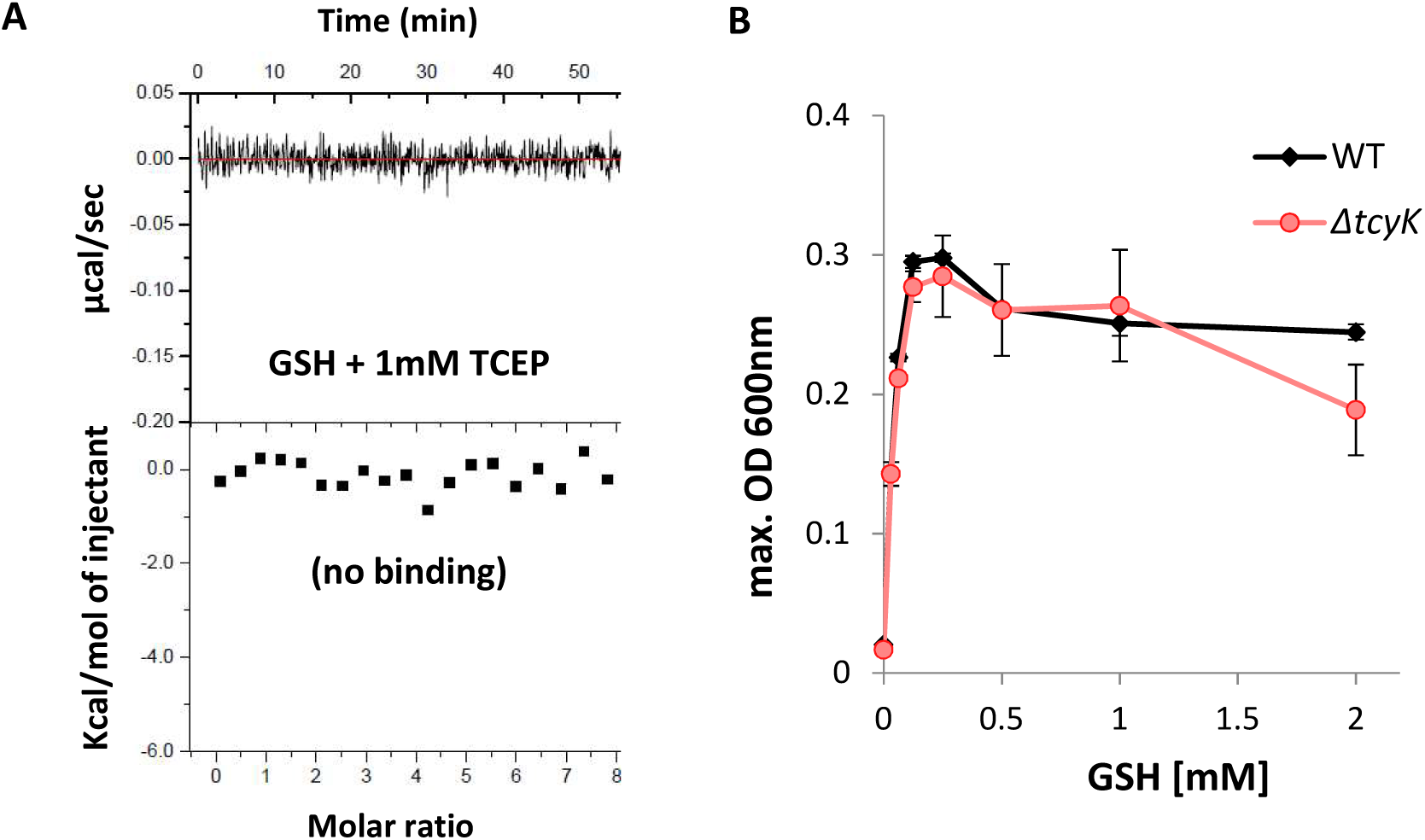
The TcyKLMN transporter does not transport glutathione. **(A)** Isothermal Titration Calorimetry (ITC) analysis, showing no binding of TcyK to reduced glutathione (GSH). Shown are the consecutive injections of 2 μL aliquots from a 200 μM GSH solution into 200 μL of 20 μM TcyK. TCEP was used as a reducing agent. The upper panels show the calorimetric titration and the lower panels display the integrated injection heat derived from the titrations. The experiment was repeated independently 3 times. **(B)** Growth of WT *Lm* and *ΔtcyK* bacteria in minimal defined media supplemented with different concentrations of reduced glutathione (GSH) as a sole source of cysteine. The data is presented as maximal OD600 values, corresponding to bacterial yield. The data represent 2 biological replicates. Error bars indicate standard deviation.

**Fig S8.**
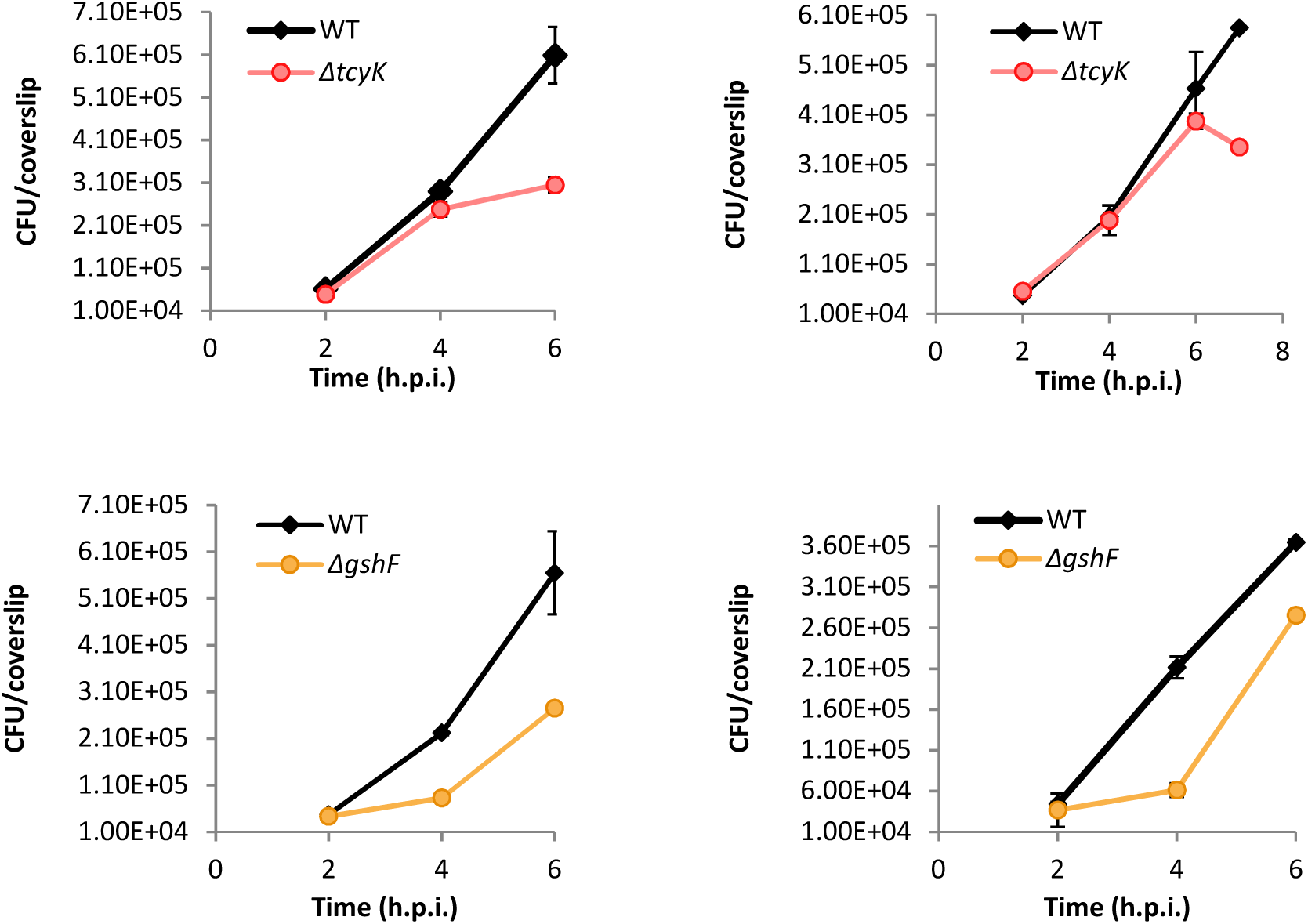
Intracellular growth of *ΔtcyK* and *ΔgshF* mutants in BMDM cells in comparison to WT *Lm*. The experiment was repeated independently 3 times, two biological repeats for each mutant are shown here, supplementary to Figure 6 C-D.

## References

1. Swaminathan B, Gerner-Smidt P. 2007. The epidemiology of human listeriosis. Microbes Infect 9:1236–43.

2. Freitag NE, Port GC, Miner MD. 2009. Listeria monocytogenes — from saprophyte to intracellular pathogen. Nat Rev Microbiol 72:623–628.

3. Lecuit M, Cossart P, Pizarro-cerda J. 2003. Invasion of mammalian cells by Listeria monocytogenes : functional mimicry to subvert cellular functions. Trends Cell Biol 13:23–31.

4. Cossart P, Vicente MF, Mengaud J, Baquero F, Perezdiaz JC, Berche P. 1989. Listeriolysin-O Is Essential for Virulence of Listeria-Monocytogenes - Direct Evidence Obtained By Gene Complementation. Infect Immun 57:3629–3636.

5. Smith G a., Marquis H, Jones S, Johnston NC, Portnoy D a., Goldfine H. 1995. The two distinct phospholipases C of Listeria monocytogenes have overlapping roles in escape from a vacuole and cell-to-cell spread. Infect Immun 63:4231–4237.

6. Marquis H, Doshi V, Portnoy DA. 1995. The broad-range phospholipase C and a metalloprotease mediate listeriolysin O-independent escape of Listeria monocytogenes from a primary vacuole in human epithelial cells. Infect Immun 63:4531–4534.

7. Grubmüller S, Schauer K, Goebel W, Fuchs TM, Eisenreich W. 2014. Analysis of carbon substrates used by Listeria monocytogenes during growth in J774A.1 macrophages suggests a bipartite intracellular metabolism. Front Cell Infect Microbiol 4:156.

8. Chico-Calero I, Suárez M, González-Zorn B, Scortti M, Slaghuis J, Goebel W, Vázquez-Boland JA. 2002. Hpt, a bacterial homolog of the microsomal glucose- 6-phosphate translocase, mediates rapid intracellular proliferation in Listeria. Proc Natl Acad Sci 99:431–436.

9. Sauer J-D, Herskovits AA, O’Riordan MXD. 2019. Metabolism of the Gram-Positive Bacterial Pathogen Listeria monocytogenes . Microbiol Spectr 7.

10. Keeney K, Colosi L, Weber W, O’Riordan M. 2009. Generation of Branched-Chain Fatty Acids through Lipoate-Dependent Metabolism Facilitates Intracellular Growth of Listeria monocytogenes. J Bacteriol 191:2187–2196.

11. Tilney LG, Portnoy DA. 1989. Actin Filaments and the Growth, Movement, and Spread of the Intacellular Bacterial Parsite, Listeria monocytogenes. JCell Biol 109:1597–1608.

12. Kocks C, Gouin E, Tabouret M, Berche P, Ohayon H, Cossart P. 1992. L. monocytogenes-induced actin assembly requires the actA gene product, a surface protein. Cell 68:521–531.

13. de las Heras A, Cain RJ, Bielecka MK, Vázquez-Boland J a. 2011. Regulation of Listeria virulence: PrfA master and commander. Curr Opin Microbiol 14:118–27.

14. Radoshevich L, Cossart P. 2017. Listeria monocytogenes : towards a complete picture of its physiology and pathogenesis. Nat Rev Microbiol 2017 161 16:32–46.

15. Lobel L, Sigal N, Borovok I, Belitsky BR, Sonenshein AL, Herskovits AA. 2015. The metabolic regulator CodY links Listeria monocytogenes metabolism to virulence by directly activating the virulence regulatory gene prfA. Mol Microbiol 95:624–644.

16. Haber A, Friedman S, Lobel L, Burg-Golani T, Sigal N, Rose J, Livnat-Levanon N, Lewinson O, Herskovits AA. 2017. L-glutamine Induces Expression of Listeria monocytogenes Virulence Genes. PLOS Pathog 13:e1006161.

17. Reniere ML, Whiteley AT, Hamilton KL, John SM, Lauer P, Brennan RG, Portnoy DA. 2015. Glutathione activates virulence gene expression of an intracellular pathogen. Nature 517:170–173.

18. Johansson J, Mandin P, Renzoni a, Chiaruttini C, Springer M, Cossart P. 2002. An RNA thermosensor controls expression of virulence genes in Listeria monocytogenes. Cell 110:551–561.

19. Ripio MT, Brehm K, Lara M, Suarez M, Vazquez-Boland JA. 1997. Glucose-1-phosphate utilization by Listeria monocytogenes is PrfA dependent and coordinately expressed with virulence factors. J Bacteriol1997/11/26. 179:7174–7180.

20. Krypotou E, Scortti M, Grundström C, Oelker M, Luisi BF, Sauer-Eriksson AE, Vázquez-Boland J. 2019. Control of Bacterial Virulence through the Peptide Signature of the Habitat. Cell Rep 26:1815–1827.e5.

21. Lobel L, Sigal N, Borovok I, Ruppin E, Herskovits AA. 2012. Integrative genomic analysis identifies isoleucine and CodY as regulators of Listeria monocytogenes virulence. PLoS Genet 8:e1002887.

22. Lobel L, Herskovits A a. 2016. Regulatory Activities of CodY Controlling Metabolism , Motility and Virulence in Listeria monocytogenes. PLOS Genet 12:1–27.

23. Levdikov VM, Blagova E, Joseph P, Sonenshein AL, Wilkinson AJ. 2006. The structure of CodY, a GTP- and isoleucine-responsive regulator of stationary phase and virulence in gram-positive bacteria. J Biol Chem 281:11366–73.

24. Sonenshein AL. 2005. CodY, a global regulator of stationary phase and virulence in Gram-positive bacteria. Curr Opin Microbiol 8:203–7.

25. Brenner M, Lobel L, Borovok I, Sigal N, Herskovits AA. 2018. Controlled branched-chain amino acids auxotrophy in Listeria monocytogenes allows isoleucine to serve as a host signal and virulence effector. PLoS Genet 14:1–21.

26. Aoyama K, Nakaki T. 2015. Glutathione in Cellular Redox Homeostasis: Association with the Excitatory Amino Acid Carrier 1 (EAAC1). Mol 2015, Vol 20, Pages 8742–8758 20:8742–8758.

27. Gopal S, Borovok I, Ofer A, Yanku M, Cohen G, Goebel W, Kreft J, Aharonowitz Y. 2005. A Multidomain Fusion Protein in Listeria monocytogenes Catalyzes the Two Primary Activities for Glutathione Biosynthesis. J Bacteriol 187:3839–3847.

28. Hall M, Grundström C, Begum A, Lindberg MJ, Sauer UH, Almqvist F, Johansson J, Sauer-Eriksson AE. 2016. Structural basis for glutathione-mediated activation of the virulence regulatory protein PrfA in Listeria. Proc Natl Acad Sci U S A 6:201614028.

29. Wang Y, Feng H, Zhu Y, Gao P. 2017. Structural insights into glutathione-mediated activation of the master regulator PrfA in Listeria monocytogenes. Protein Cell 8:308–312.

30. Tsai HN, Hodgson DA. 2003. Development of a Synthetic Minimal Medium for Listeria monocytogenes. Appl Environ Microbiol 69:6943–6945.

31. Xayarath B, Marquis H, Port GC, Freitag NE. 2009. Listeria monocytogenes CtaP is a multifunctional cysteine transport-associated protein required for bacterial pathogenesis. Mol Microbiol 74:956–973.

32. Schauer K, Geginat G, Liang C, Goebel W, Dandekar T, Fuchs TM. 2010. Deciphering the intracellular metabolism of Listeria monocytogenes by mutant screening and modelling. BMC Genomics 11:573.

33. Friedman S, Linsky M, Lobel L, Rabinovich L, Sigal N, Herskovits AA. 2017. Metabolic genetic screens reveal multidimensional regulation of virulence gene expression in Listeria monocytogenes, and an aminopeptidase that is critical for PrfA protein activation. Infect Immun IAI.00027–17.

34. Burguière P, Auger S, Hullo M, Danchin A, Martin-verstraete I, Burguie P. 2004. Three Different Systems Participate in l-Cystine Uptake in Bacillus subtilis Three Different Systems Participate in L -Cystine Uptake in Bacillus subtilis. J Bacteriol 186:4875–4884.

35. Tanous C, Soutourina O, Raynal B, Hullo MF, Mervelet P, Gilles AM, Noirot P, Danchin A, England P, Martin-Verstraete I. 2008. The CymR regulator in complex with the enzyme CysK controls cysteine metabolism in Bacillus subtilis. J Biol Chem 283:35551–35560.

36. Burguière P, Fert J, Guillouard I, Danchin A, Martin-verstraete I, Burguie P, Auger S. 2005. Regulation of the Bacillus subtilis ytmI Operon, Involved in Sulfur Metabolism. J Bacteriol 187:6019–6030.

37. CM C, A D, P M, A S. 2014. Paralogous metabolism: S-alkyl-cysteine degradation in Bacillus subtilis. Environ Microbiol 16:101–117.

38. Even S, Burguière P, Auger S, Danchin A, Martin-verstraete I, Burguie P, Soutourina O. 2006. Global Control of Cysteine Metabolism by CymR in Bacillus subtilis. J Bacteriol 188:2184–2197.

39. Rees DC, Johnson E, Lewinson O. 2009. ABC transporters: The power to change. Nat Rev Mol Cell Biol 10:218–227.

40. Biswas R, Sonenshein AL, Belitsky BR. 2020. Role of GlnR in Controlling Expression of Nitrogen Metabolism Genes in Listeria monocytogenes. J Bacteriol 202:209–229.

41. Marquis H, Bouwer HGA, Hinrichs DJ, Portnoy DA. 1993. Intracytoplasmic growth and virulence of Listeria monocytogenes auxotrophic mutants. Infect Immun 61:3756–3760.

42. Reniere ML, Whiteley AT, Portnoy DA. 2016. An In Vivo Selection Identifies Listeria monocytogenes Genes Required to Sense the Intracellular Environment and Activate Virulence Factor Expression. PLoS Pathog 12:1–27.

43. Ji Q, Zhang L, Sun F, Deng X, Liang H, Bae T, He C. 2012. Staphylococcus aureus CymR is a new thiol-based oxidation-sensing regulator of stress resistance and oxidative response. J Biol Chem 287:21102–21109.

44. Virginie M, Nakaura Y, P SR, Yamaguchi H, Losick R, Fujita Y, Sonenshein AL. 2003. Additional targets of the Bacillus subtilis global regulator CodY identified by chromatin immunoprecipitation and genome-wide transcript analysis. J Bacteriol 185:1911–1922.

45. Brinsmade SR, Alexander EL, Livny J, Stettner AI, Segrè D, Rhee KY, Sonenshein AL. 2014. Hierarchical expression of genes controlled by the Bacillus subtilis global regulatory protein CodY. Proc Natl Acad Sci 111:8227–8232.

46. Combs JA, DeNicola GM. 2019. The Non-Essential Amino Acid Cysteine Becomes Essential for Tumor Proliferation and Survival. Cancers (Basel) 11.

47. Sato H, Tamba M, Ishii T, Bannai S. 1999. Cloning and Expression of a Plasma Membrane Cystine/Glutamate Exchange Transporter Composed of Two Distinct Proteins *. J Biol Chem 274:11455–11458.

48. Kang M, Eichhorn CD, Feigon J. 2014. Structural determinants for ligand capture by a class II preQ1 riboswitch. Proc Natl Acad Sci U S A 111:E663–71.

49. Shih AY, Erb H, Sun X, Toda S, Kalivas PW, Murphy TH. 2006. Cystine/Glutamate Exchange Modulates Glutathione Supply for Neuroprotection from Oxidative Stress and Cell Proliferation. J Neurosci 26:10514–10523.

50. Kortebi M, 1☯ M, Mitchell G, Choux CP, Prevost M-C, Cossart P, Bierne H. 2017. Listeria monocytogenes switches from dissemination to persistence by adopting a vacuolar lifestyle in epithelial cells https://doi.org/10.1371/journal.ppat.1006734.

51. Birmingham CL, Canadien V, Kaniuk NA, Steinberg BE, Higgins DE, Brumell JH. 2008. Listeriolysin O allows Listeria monocytogenes replication in macrophage vacuoles. Nat 2008 4517176 451:350–354.

52. Wong J, Chen Y, Gan YH. 2015. Host cytosolic glutathione sensing by a membrane histidine kinase activates the type VI secretion system in an intracellular bacterium. Cell Host Microbe 18:38–48.

53. Alkhuder K, Meibom KL, Dubail I, Dupuis M, Charbit A. 2009. Glutathione provides a source of cysteine essential for intracellular multiplication of Francisella tularensis. PLoS Pathog 5.

54. Portman JL, Dubinsky SB, Peterson BN, Whiteley AT, Portnoy DA. 2017. Activation of the Listeria monocytogenes Virulence Program by a Reducing Environment. MBio 8:1–13.

55. Vergauwen B, Elegheert J, Dansercoer A, Devreese B, Savvides SN. 2010. Glutathione import in Haemophilus influenzae Rd is primed by the periplasmic heme-binding protein HbpA. Proc Natl Acad Sci 107:13270–13275.

56. Vergauwen B, Verstraete K, Senadheera DB, Dansercoer A, Cvitkovitch DG, Guédon E, Savvides SN. 2013. Molecular and structural basis of glutathione import in Gram-positive bacteria via GshT and the cystine ABC importer TcyBC of Streptococcus mutans. Mol Microbiol 89:288–303.

57. Simon R, Priefer U, Pühler A. 1983. A Broad Host Range Mobilization System for In Vivo Genetic Engineering: Transposon Mutagenesis in Gram Negative Bacteria. Nat Biotechnol 1:784–791.

58. Phan-thanh L, Gormon T. 1997. A chemically defined minimal medium for the optimal culture of Listeria. Int J Food Microbiol 35:91–95.

59. Bron P a, Monk IR, Corr SC, Hill C, Gahan CGM, Corr C. 2006. Novel Luciferase Reporter System for In Vitro and Organ-Specific Monitoring of Differential Gene Expression in Listeria monocytogenes. Appl Environ Microbiol 72:2876–2884.

60. Stead MB, Agrawal A, Bowden KE, Nasir R, Mohanty BK, Meagher RB, Kushner SR. 2012. RNAsnap™: a rapid, quantitative and inexpensive, method for isolating total RNA from bacteria. Nucleic Acids Res 40:e156.

61. Bailey TL, Boden M, Buske FA, Frith M, Grant CE, Clementi L, Ren J, Li WW, Noble WS. 2009. MEME Suite: Tools for motif discovery and searching. Nucleic Acids Res 37:202–208.

62. Levdikov VM, Blagova E, Young VL, Belitsky BR, Lebedev A, Sonenshein AL, Wilkinson AJ. 2017. Structure of the branched-chain Amino Acid and GTP-sensing global regulator, CodY, from Bacillus subtilis. J Biol Chem 292:2714–2728.

63. Solovyev V, Salamov A. 2010. Automatic Annotation of Bacterial Community Sequences and Application To Infections Diagnostic. Metagenomics its Appl Agric Biomed Environ Stud 61–78.

64. Wurtzel O, Sesto N, Mellin JR, Karunker I, Edelheit S, Bécavin C, Archambaud C, Cossart P, Sorek R. 2012. Comparative transcriptomics of pathogenic and non-pathogenic Listeria species. Mol Syst Biol 8:583.

65. Crimmins GT, Herskovits AA, Rehder K, Sivick KE, Lauer P, Dubensky TW, Portnoy D a. 2008. Listeria monocytogenes multidrug resistance transporters activate a cytosolic surveillance pathway of innate immunity. Proc Natl Acad Sci U S A 105:10191–6.

66. Sigal N, Pasechnek A, Herskovits AA. 2016. RNA purification from intracellularly grown listeria monocytogenes in macrophage cells. J Vis Exp 2016:1–7.

